# Emergence of RNA-guided transcription factors via domestication of transposon-encoded TnpB nucleases

**DOI:** 10.1101/2023.11.30.569447

**Authors:** Tanner Wiegand, Florian T. Hoffmann, Matt W.G. Walker, Stephen Tang, Egill Richard, Hoang C. Le, Chance Meers, Samuel H. Sternberg

## Abstract

Transposon-encoded *tnpB* genes encode RNA-guided DNA nucleases that promote their own selfish spread through targeted DNA cleavage and homologous recombination^1–4^. This widespread gene family was repeatedly domesticated over evolutionary timescales, leading to the emergence of diverse CRISPR-associated nucleases including Cas9 and Cas12^5,6^. We set out to test the hypothesis that TnpB nucleases may have also been repurposed for novel, unexpected functions other than CRIS-PR-Cas. Here, using phylogenetics, structural predictions, comparative genomics, and functional assays, we uncover multiple instances of programmable transcription factors that we name TnpB-like nuclease-dead repressors (TldR). These proteins employ naturally occurring guide RNAs to specifically target conserved promoter regions of the genome, leading to potent gene repression in a mechanism akin to CRISPRi technologies invented by humans^7^. Focusing on a TldR clade found broadly in *Enterobacteriaceae*, we discover that bacteriophages exploit the combined action of TldR and an adjacently encoded phage gene to alter the expression and composition of the host flagellar assembly, a transformation with the potential to impact motility^8^, phage susceptibility^9^, and host immunity^10^. Collectively, this work showcases the diverse molecular innovations that were enabled through repeated exaptation of genes encoded by transposable elements, and reveals that RNA-guided transcription factors emerged long before the development of dCas9-based editors.

---

Transposons play a central role in driving genome evolution and genome expansion, due to their proliferative nature and capacity for horizontal gene transfer, and the genes responsible for their mobility are among the most abundant genes in all of nature^11^. Although transposons are intrinsically selfish, having evolved strategies to increase their likelihood of survival without necessarily benefitting the host, their molecular adaptations have directly shaped, and been shaped by, co-evolving processes in the organisms that maintain them^12^. Thus, whereas unchecked transposition poses a perennial cellular threat that has spurred the evolution of cellular defense mechanisms, transposons also encode a vast repertoire of diverse enzymes that have been repeatedly repurposed by hosts, leading to the emergence of biological pathways as varied as intron splicing, immunoglobulin gene diversification, genome rearrangement, and genome defense. Indeed, some of the most intricate cellular reactions involving nucleic acids have arisen in the genetic conflict, cooperation, and cooption between cells and transposable elements.

The origins of bacterial adaptive immune systems known as CRISPR-Cas can be traced directly to such host-transposon interactions, whereby enzymes originally adapted for transposition were exapted^13^ for novel roles in viral DNA acquisition and targeting^14,15^. The ‘universal’ *cas1* gene encodes an integrase responsible for preserving memories of past infections by splicing viral DNA fragments into the CRISPR array, in a biochemical reaction reminiscent of transposon integration^16,17^, and ancestral CRISPR-less Cas1 homologs perform similar reactions within the context of transposable elements known as casposons^18,19^. Analogously, recent studies have demonstrated that the biochemical activities of Cas9 and Cas12, which perform RNA-guided DNA binding and cleavage during an immune response, can be traced back to ancestral transposon-encoded nucleases known as IscB and TnpB, respectively^1,2^, which perform similar reactions to promote transposon maintenance and spread^3,4^. In turn, nuclease-deficient CRISPR-Cas systems have been repurposed by transposons on at least four independent occasions, to facilitate a novel RNA-guided DNA integration pathway mediated by CRIS-PR-associated transposases (CASTs) ^20–22^. CRISPR-Cas evolution and diversification can therefore be considered as having arisen through recurrent gene domestication events that leverage the fundamental molecular properties of the proteins or protein-RNA complexes being acquired. Similar cooption events have also frequently occurred between bacteria and bacteriophages^23,24^, highlighting the extensive flux of genetic information between hosts and diverse types of mobile genetic elements (MGEs) ^25^.

Transposon-encoded TnpB proteins represent a vast reservoir of RNA-guided nucleases that are found in association with diverse transposons/transposases across all three domains of life^26–28^. In bacteria, *tnpB* genes are encoded within IS200/IS605- and IS607-family transposons, which are minimal selfish genetic elements that are mobilized by a TnpA-family transposase but often exist in a non-autonomous form^29^. These transposons harbor conserved left end (LE) and right end (RE) sequences that define the boundaries of the mobile DNA, and in addition to protein-coding genes, they also encode non-coding RNAs, referred to as ωRNA (or reRNA), that feature a scaffold region spanning the transposon RE and a ∼16-nt guide derived from the transposon-flanking sequence^1,2^ (**Fig. 1a**). We recently demonstrated that TnpA-mediated transposition generates a scarless excision product at the donor site that is rapidly recognized and cleaved by TnpB-ωRNA complexes, in a reaction dependent on RNA-DNA complementarity and the presence of a cognate transposon/target-adjacent motif (TAM), leading to transposon reinstallation via DSB-mediated homologous recombination^3,4^. RNA-guided DNA cleavage thus first arose as a biochemical strategy to prevent permanent transposon loss and was only later adapted for a role in antiviral immunity.

**Figure 1.**
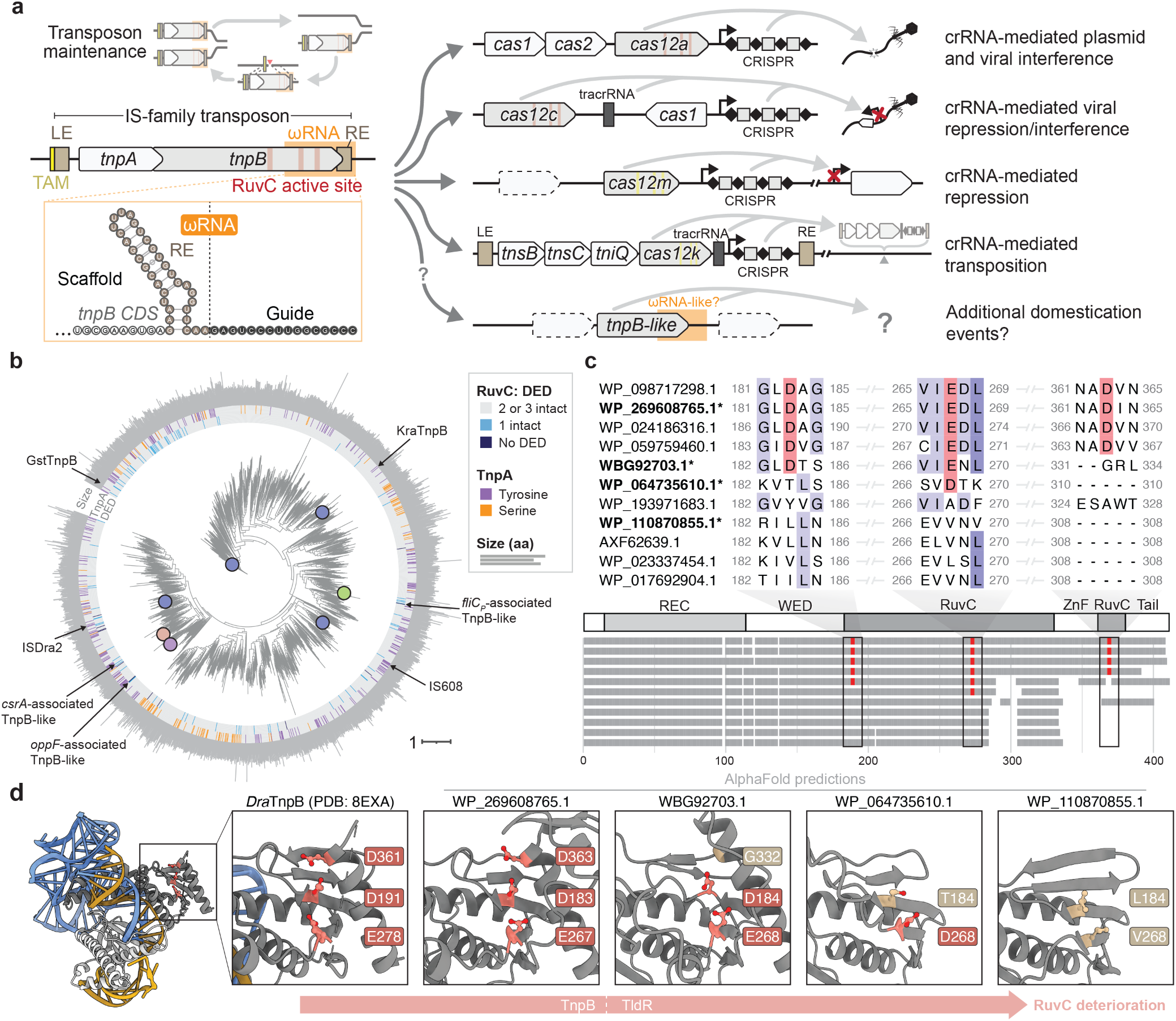
Bioinformatic identification of naturally occurring, nuclease-deficient TnpB homologs. **a,** Canonical TnpB proteins are encoded by bacterial transposons known as IS elements, and exhibit RNA-guided nuclease activity that maintains transposons at sites of excision during transposition (left). Domestication of *tnpB* genes led to the evolution of diverse CRISPR-associated *cas12* derivatives, with diverse functions and mechanisms (right). LE, transposon left end; RE, right end; ωRNA, transposon-encoded guide RNA; crRNA, CRISPR RNA. **b,** Phylogenetic tree of TnpB proteins, with previously studied homologs and newly identified TnpB-like nuclease-dead repressor (TldR) proteins highlighted. The rings indicate RuvC DED active site intactness (inner), TnpA transposase association (middle) and protein size (outer). **c,** Multiple sequence alignment of representative TnpB and TldR sequences, highlighting deterioration of RuvC active site motifs and loss of the C-terminal Zinc-finger (ZnF)/RuvC domain. **d,** Empirical (DraTnpB) and predicted AlphaFold structures of TnpB and TldR homologs marked with an asterisk in **c,** showing progressive loss of the active site catalytic triad.

TnpB nucleases have been independently domesticated numerous times over evolutionary timescales, leading to the emergence of dozens of unique CRISPR-Cas12 subtypes that feature diverse guide RNA requirements and PAM specificities^30,31^. In nearly all cases, Cas12 homologs rely on the same RuvC nuclease domain as TnpB for target cleavage, highlighting its conserved role in nucleic acid chemistry. However, recent studies uncovered atypical Cas12 homologs — Cas12c and Cas12m — that have lost the ability to cleave target DNA but instead bind and repress gene transcription as an alternative mechanism to preventing MGE proliferation^32,33^. Type V-K CASTs similarly rely on nuclease-inactivated Cas12k homologs that are still active for RNA-guided DNA binding, leaded to programmable transposition^21,22,34^ (**Fig. 1a**). Given the sheer abundance of *tnpB* genes and the pro-found utility of RNA-guided DNA binding — as exemplified both in biology and biotechnology^35^ — we hypothesized that TnpB-like proteins may have been domesticated for novel functions, and we set out to test this hypothesis by specifically mining for nuclease-inactivated variants located in diverse genetic neighborhoods.

Here we report the discovery of a novel family of TnpB-like nuclease-dead repressors (hereafter TldR) that function not for transposition, but for RNA-guided transcriptional control, thus rendering the name “TnpB (transposase B)” inapposite. Using a custom bioinformatics pipeline, we identified multiple independent TldR clades that evolved from transposon-encoded TnpB nucleases via RuvC active site deterioration, coincident with newly acquired, non-transposase gene associations. TldRs function with adjacently encoded non-coding guide RNAs (gRNAs) to target complementary DNA sequences flanked by a TAM within promoter regions, and target binding down-regulates gene expression through competitive exclusion of RNA polymerase. Remarkably, flagellin (FliC)-associated TldR homologs are exploited by prophages to specifically remodel the host flagellar apparatus, which we discovered using *in vivo* genetic perturbation experiments in a clinical *Enterobacter* isolate. Collectively, this work reveals a novel evolutionary trajectory of transposon-derived, RNA-guided nucleases, and highlights the molecular opportunities afforded by transposon gene exaptation.

## RESULTS

### Bioinformatic identification of nuclease-dead TnpB proteins

We developed a bioinformatics pipeline to identify TnpB proteins with inactivating mutations in the RuvC domain, motivated by the hypothesis that these would represent likely gene exaptations for functions beyond transposon proliferation. We clustered a multiple sequence alignment of 95,731 unique TnpB-like sequences at 50% sequence identity and then performed an automatic assessment of the conservation of RuvC active site residues. TnpB, like Cas12 nucleases, harbors a catalytic motif consisting of three acidic residues (DED), and mutating any residue in this motif abolishes nuclease activity^2,3,36^. However, recent analyses of TnpBs and eukaryotic TnpB-like proteins (i.e., Fanzors) indicate that one of the catalytic residues can occur at an alternate position in the RuvC domain^27^. Indeed, we observed that this flexibility often resulted in the spurious identification of catalytically inactivated TnpB-like proteins, since structural predictions and manual inspections suggested an intact catalytic triad. Thus, we restricted our initial analysis to TnpB-like proteins with two or more mutations in the RuvC DED motif.

This search, supplemented with additional homologs identified in more focused analyses described below, identified 506 unique TnpB-like proteins with conserved mutations that are predicted to inactivate the RuvC nuclease domain (**Fig. 1b, Supplementary Dataset 1**). The polyphyletic distribution of these inactivated nucleases suggest that they emerged on multiple occasions independently (**Fig. 1b**), and based on their predicted role in transcriptional repression (see below), we refer to them hereafter as TnpB-like nuclease-dead repressors (TldRs). Interestingly, TldRs exhibit a range of deteriorated active sites, with one, two or all three acidic residues mutated, and many homologs also feature truncated C-terminal domains that ablate RuvC and zinc-finger (ZnF) domains (**Fig. 1c, Extended Data Fig. 1**). AlphaFold predictions provided further structural support for the sequential deterioration of the RuvC active site, without any apparent degradation in the remainder of the overall TnpB/TldR fold or the RNA binding interface (**Fig. 1c**), suggesting the intriguing possibility that RNA-guided DNA targeting functions could be preserved for these inactivated nucleases.

### *tldRs* associate with novel genes and are mobilized by temperate phages

Canonical *tnpB* genes in bacteria, alongside their ωRNA guides, are encoded within IS200/ IS605- or IS607-family transposons that can be straightforwardly identified using both comparative genomics and by defining the transposon left end (LE) and right end (RE) ^1–4^; in addition, a hallmark feature is their frequent association with *tnpA* transposase genes^29,37^ (**Fig. 2a, left**). Remarkably, the genomic context surrounding *tldR* genes consistently lacked *tnpA* and identifiable LE/RE sequences, and instead, we observed strong genetic associations with novel, non-transposon genes that were clade specific (**Fig. 1b, 2a**). One TldR group is consistently associated with five to six genes encoding components of ABC transporter systems^38,39^, the last of which is *oppF*, and is mainly present in *Enterococci* genomes (**Supplementary Dataset 2**). A second TldR group is tightly associated with *fliC*, a gene encoding the flagellin subunit of flagellar assemblies that propel bacteria in aqueous environments^10^, and is found in diverse *Enterobacteriaceae* (**Supplementary Dataset 3**). A third TldR group from Clostridial genomes is similarly associated with flagellin genes, in addition to a carbon storage regulator (*csrA*) that is involved in flagellar subunit regulation^40^ (**Supplementary Dataset 1**). In all three cases, we observed loci encoding TldRs and their associated genes in varied genetic contexts, suggesting that they have maintained their associations over long time scales and/or that they have been mobilized in tandem. Strong genetic associations are also often indicative of functional coupling^41^, indicating that

**Figure 2.**
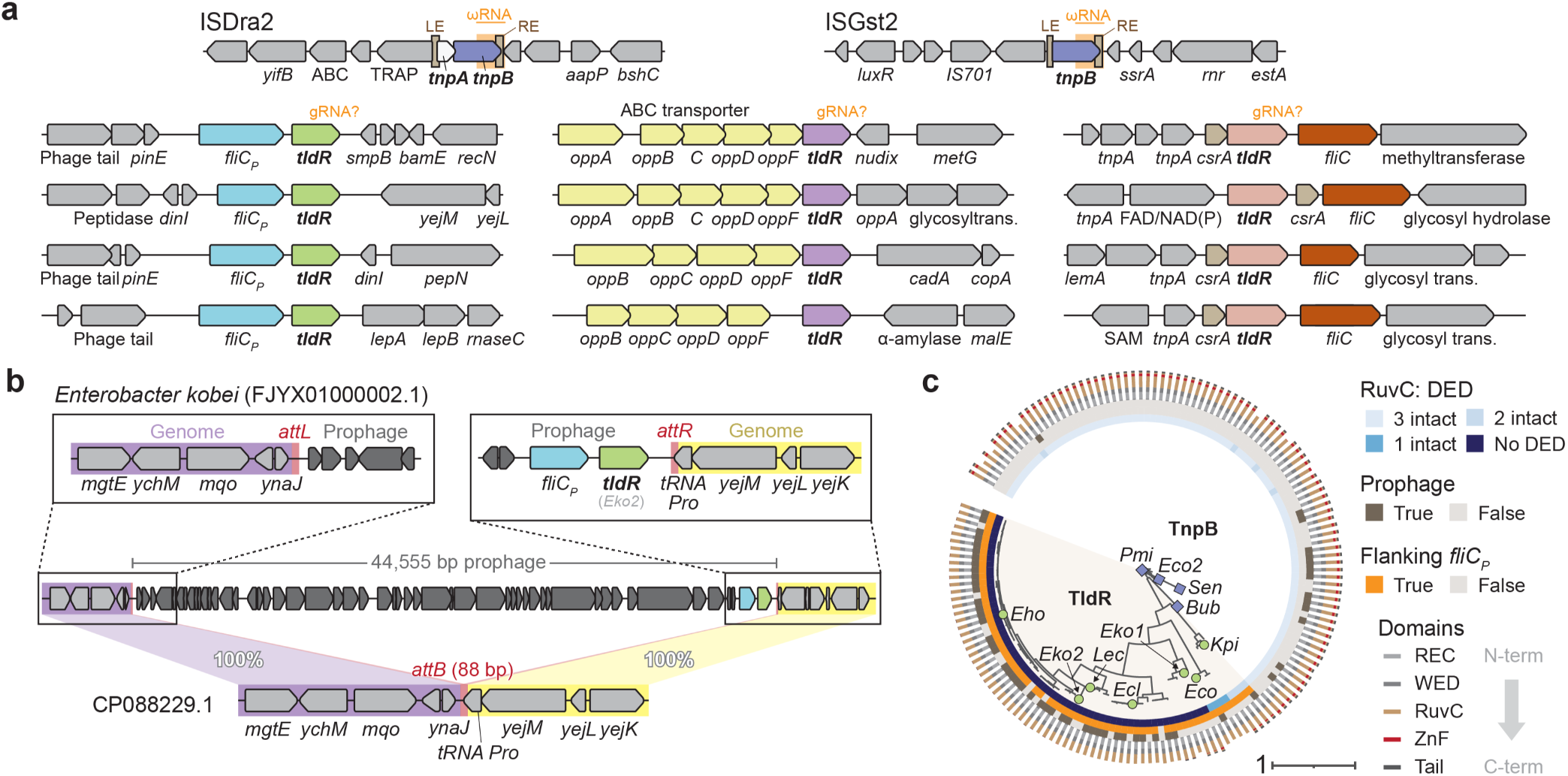
*tldR* genes are strongly associated with diverse non-transposon genes and encoded in prophages. **a,** Genomic architecture of well-studied transposons that encode TnpB (top), and of novel regions that encode TldR proteins (bottom) in association with prophage-encoded *fliC_P_* (left)*, oppF* and ABC transporter operons (middle), and a transcriptional regulator (*csrA*) of an accompanying *fliC* (right). **b,** Comparison of a representative *fliC_P_-tldR* locus with a closely related *Enterobacter kobei* strain reveals that the entire locus is encoded within the boundaries of the prophage element, with identifiable recombination sequences (*attL/attR/attB*). **c,** Phylogenetic tree of *fliC_P_-*associated TldR proteins from **a**, together with closely related TnpB proteins that contain intact RuvC active sites. The rings indicate RuvC DED active site intactness (inner), prophage association (middle), *fliC_P_* association (middle), and TldR/TnpB domain composition (outer). Prophage association was defined as true if the homolog was encoded within 20 kbp of five or more genes with a phage annotation; *fliC_P_* association was defined as true if the homolog was encoded within three ORFs of a *fliC* homolog. Homologs marked with a blue square (TnpB) or green circle (TldR) were tested in heterologous experiments.

TldR proteins may be involved in flagellar and ABC transporter expression and/or assembly pathways. A closer inspection of genomic loci encoding *fliC-tldR* revealed the striking presence of numerous upstream genes with bacteriophage (phage) annotations, suggesting the potential presence of an integrated prophage (**Fig. 2a, Supplementary Data Fig. 1a**). When we used BLAST to search the NCBI non-redundant and whole genome shotgun databases, we identified genomes that were highly similar to those encoding *fliC-tldR* but lacked phage genes, enabling us to confidently annotate the prophage boundaries and conserved *attL*/*attR* recombination sequences (**Fig. 2b, Extended Data Fig. 2a**). These analyses indicate that both *tldR* and its associated phage-encoded *fliC* (hereafter *fliC_P_*) are core components of temperate phage genomes, suggesting the possibility of a novel role in promoting viral infection or lysogenization. Consistent with this hypothesis, the genetic association between *tldR and fliC_P_* emerged coincident with the acquisition of nuclease-inactivating mutations in the RuvC domain (**Fig. 2c**).

To further establish the robustness of these conclusions, we analyzed additional prophage elements and found that *fliC_P_-tldR* loci are encoded within temperate phages that, in some cases, share less than ten percent genomic sequence conservation (**Extended Data Fig. 2b,c**). Additional BLAST searches revealed two metagenome-assembled phage genomes in the taxa *Caudovirales* that encode *fliC_P_-tldR* (**Supplementary Data Fig. 1b**). Collectively, these data demonstrate that at least one TnpB domestication event involved the loss of nuclease activity, the loss of flanking transposon end sequences, and the gain of an accessory gene possibly linked to a novel function in phage biology. Notably, no similar bacteriophage associations were detected for *oppF*- or *csrA-*associated TldRs.

### Identification of TldR-associated guide RNAs that target conserved promoters

We reasoned that identifying the putative guide RNA substrates of TldRs would provide a major clue to their biological function, and thus made this our next objective. Transposon-encoded TnpB proteins function together with gRNAs (also referred to as reRNAs) that are transcribed from within or near the 3′-end of the *tnpB* coding sequence, to perform RNA-guided DNA cleavage^1,2^. Like CRISPR RNAs, gRNAs harbor both an invariant ‘scaffold’ sequence that is a binding site for TnpB, as well as the ‘guide’ sequence that specifies target sites through complementary RNA-DNA base-pairing. Importantly, the gRNA sequence extending beyond the transposon right end (RE) invariably comprises the guide for TnpB, and numerous *in silico* strategies can therefore be applied for gRNA identification, including comparative genomics, the ISfinder database^42^, covariance models of the gRNA structure, and sequence alignments (**Fig. 3a**). Using these strategies, we identified the LE/RE boundaries and gRNAs associated with nuclease-active TnpB homologs that are closely related to *fliC_P_* and *oppF*-associated TldRs (**Fig. 3b**). Similar analyses also revealed the predicted 3–5-bp transposon/target-adjacent motif (TAM) sequences recognized by these TnpB homologs during DNA binding and cleavage^1–3^ **(Fig. 3b**), akin to the role of PAM in DNA binding and cleavage by CRISPR-Cas9 and Cas12^43^.

**Figure 3.**
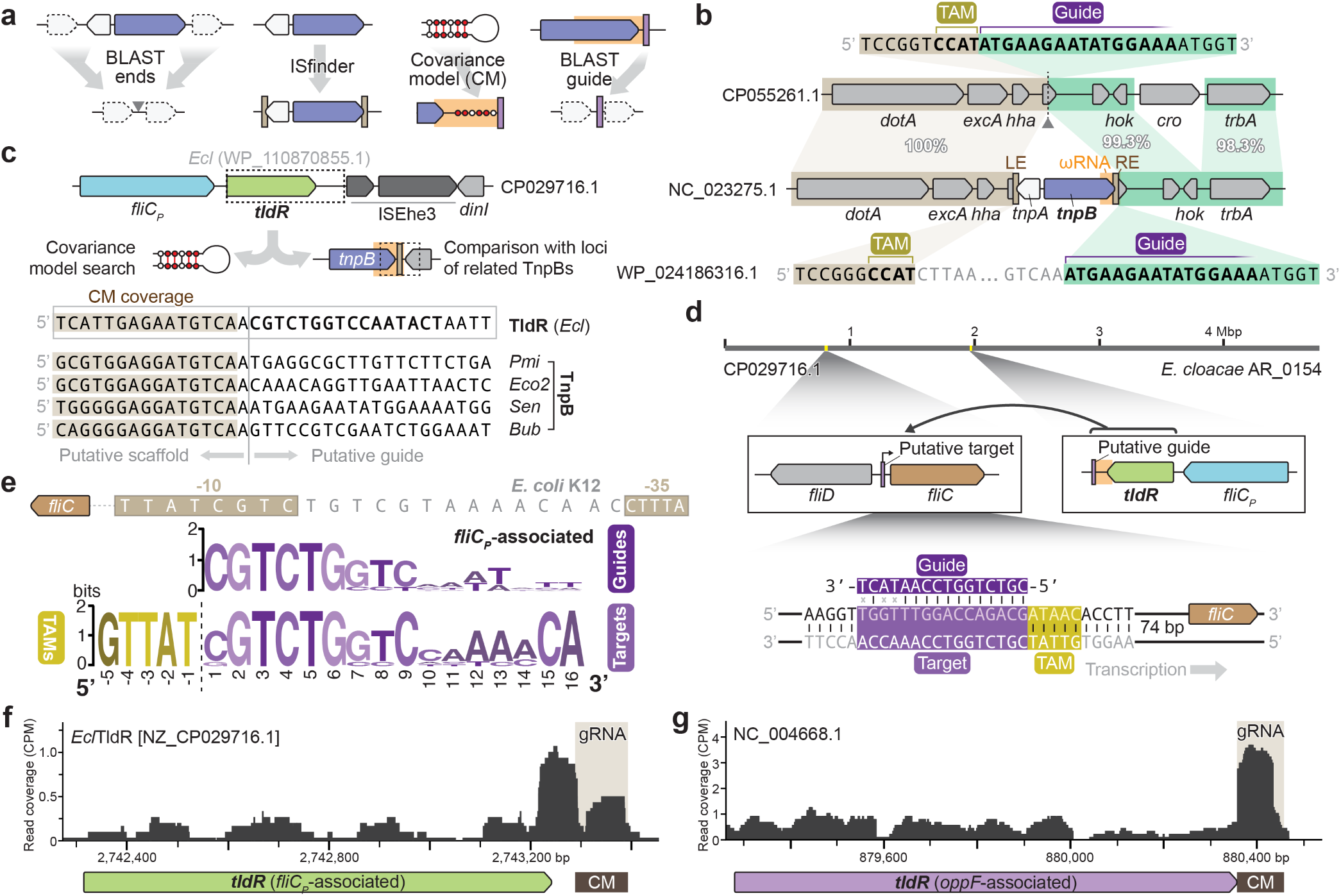
TldR proteins are encoded next to gRNAs that target conserved genomic sites. **a,** Bioinformatic strategies to investigate *tldR*/*tnpB* loci, including comparative genomics, searching within the ISfinder database, gRNA prediction using covariance models, and target prediction using BLAST. **b,** Representative *tnpB* locus and an isogenic locus above that lacks the IS element. Comparison of both sequences reveals the putative TAM recognized by TnpB, which flanks the transposon LE, and the guide portion of the ωRNA, which flanks the transposon RE. **c,** Schematic of a representative *fliC_P_- tldR* locus from *Enterobacter cloacae* (top), and bioinformatics approach to predict the gRNA sequence using both CM search and comparison to related *tnpB* loci. This analysis identified the putative scaffold and guide portions of TldR- and TnpB-associated gRNAs (bottom). **d,** Analysis of the guide sequence from the *Ecl*TldR-associated gRNA in **c** revealed a putative genomic target near the predicted promoter of a distinct (host) copy of *fliC* located ∼1 Mbp away (middle). The magnified schematic at the bottom shows the predicted TAM and gRNA-target DNA base-pairing interactions relative to the *fliC* start codon. **e**, Annotated −10 and −35 promoter elements upstream of *fliC* recognized by FliA/σ^28^ in *E. coli* K12 (top), and WebLogos of predicted guides and genomic targets associated with diverse *fliC_P_*-associated TldRs from Fig. 2c (bottom). **f**-**g**, Published RNA-seq data for *Enterobacter cloacae* (**f**) and *Enterococcus faecalis* (**g**) reveal evidence of native *tldR* and gRNA expression for *fliC_P_*- and *oppF*-associated TldRs, respectively. The predicted gRNAs from CM analyses are indicated; unique genome-mapping reads are shown as overlays of three replicates.

The absence of identifiable transposon ends flanking *tldR* rendered similar annotations of its guide RNA unfeasible, so we used covariance models (CM) built from gRNA sequences of related TnpBs. We hypothesized that TldR-associated guide RNAs (gRNA) would be encoded near the gene, and after scanning a 500-bp window flanking each *tldR* gene with the gRNA CM, we identified high-confidence gRNA-like sequences for both *fliC_P_*- and *oppF*-associated *tldR* loci (**Supplementary Data Fig. 2**). In both cases, these RNAs were encoded downstream of *tldR*, similar to other *tnpB-*gRNA loci, suggesting that functional interactions with a guide RNA may have been preserved in the face of nuclease-inactivating mutations. The strong conservation at the 3′ end of the gRNA scaffold allowed us to further predict the junction between the scaffold and putative guide sequence (**Fig. 3c, Supplementary Data Fig. 2**).

Using these putative guide sequences as queries, we next performed BLAST searches to identify potential genomic targets of *fliC_P_*-associated TldR. The strongest match was in a genomic region that encodes other flagellar components, and strikingly, was specifically located in the intergenic region between *fliD* and a second (host) *fliC* gene distinct from the prophage-encoded *fliC_P_* ortholog (**Fig. 3d**). In *E. coli*, *fliC* expression is regulated by an alternative sigma factor (σ^28^) also known as FliA^44,45^, and the putative target of the TldR-associated gRNA directly overlapped the FliA –10 promoter element, and was flanked by a conserved GTTAT motif that is highly similar to the TAM recognized by TnpB nucleases similar to TldR (**Fig. 3e**). Remarkably, these sequence features — similarity between the putative gRNA guide and *fliC* promoter, abutted by a cognate TAM — were strongly conserved across all *fliC_P_*-associated loci analyzed.

Having identified putative gRNA sequences and prospective TldRs guide-target pairs, we next sought evidence for native *tldR* and gRNA expression. When we analyzed RNA sequencing datasets^46^ from organisms with *fliC_P_-tldR* or *oppF*-*tldR* that are available on the NCBI short read archive (SRA) and gene expression omnibus (GEO), we observed read coverage over the regions identified by our CM search (**Fig. 3f,g**), providing additional evidence of functional gRNA expression from regions flanking *tldR* loci.

Collectively, these observations indicated that nuclease-inactivated *tnpB* genes remain strongly associated with noncoding RNA loci, and they immediately suggested a model for *fliC_P_*-*tldR* function, wherein phage-encoded TldR-gRNA complexes could repress expression of the host FliC protein while producing their own FliC_P_ homolog. Notably, the substantial sequence differences between host and prophage-encoded FliC and FliC_P_ homologs, specifically within the hypervariable central domains, revealed the potential biological implications of this organellar transformation (see below).

### RIP-seq reveals mature gRNA substrates and putative OppF-TldR targets

To determine if TldR proteins bind their associated guide RNAs, we cloned a representative FLAG-tagged *fliC_P_*-associated TldR (*Eho*TldR) and *oppF*-associated TldR (*Efa1*TldR) into expression plasmids, alongside 240 bp encompassing the putative gRNA scaffold and a 20-bp guide sequence (**Fig. 4a**). After performing RNA immunoprecipitation sequencing (RIP-seq) and mapping reads to the *E. coli* genome and expression plasmid, we identified a mature, ∼113-nt gRNA for *Eho*TldR that encompassed a 97-nt scaffold upstream of a 16-nt guide, indicating processing from the initial transcript down to a final mature form (**Fig. 4a**). Previous work has shown that TnpB proteins catalyze processing of their own guides through a RuvC-dependent mechanism^4,47^, however the absence of an intact catalytic triad in TldR proteins suggests that the mature gRNA may instead represent the sequence protected from cleavage by cellular ribonucleases.

**Figure 4.**
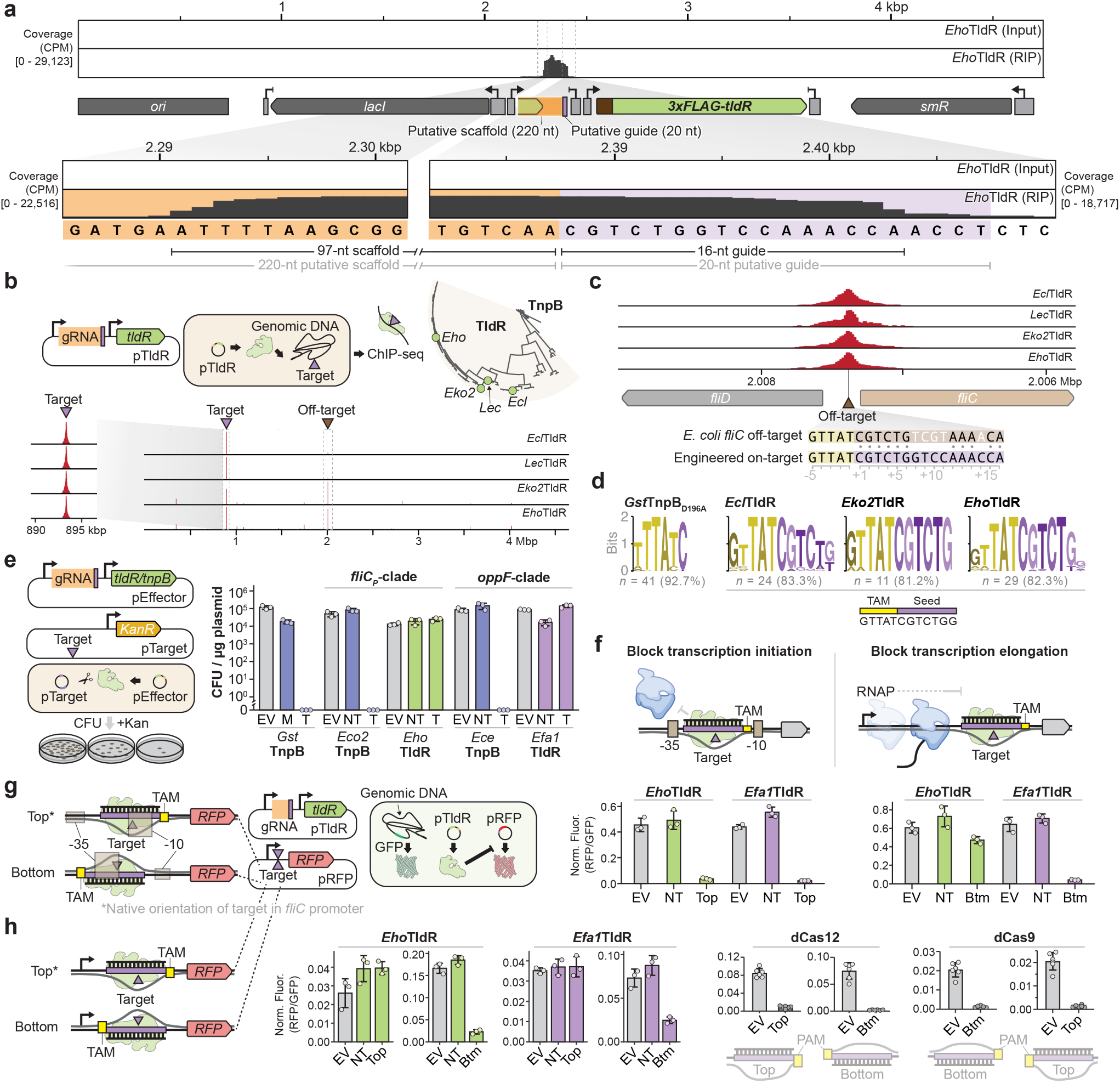
TldRs are RNA-guided DNA-binding proteins capable of programmable transcriptional repression. **a,** RNA immunoprecipitation sequencing (RIP-seq) data from a *fliC_P_-*associated TldR homolog from *Enterobacter hormaechei* (*Eho*TldR) reveals the boundaries of a mature gRNA containing a 16-nt guide sequence. Reads were mapped to the TldR-gRNA expression plasmid; an input control is shown. **b,** Schematic of chromatin immunoprecipitation DNA sequencing (ChIP-seq) approach to investigate RNA-guided DNA binding for TldR candidates (top), and representative ChIP-seq data for four homologs revealing strong enrichment at the expected genomic target site and a prominent off-target (bottom). **c,** Magnified view of ChIP-seq peaks at the labeled off-target site in **b**, which corresponds to a TAM and partially matching target sequence at the promoter of *E. coli* K12 *fliC.* **d,** Analysis of conserved motifs bound by the indicated TldR homolog using MEME ChIP, which reveals specificity for the TAM and a ∼6-nt seed sequence. The number of peaks and percentage of total called peaks contributing to each motif is indicated; low occupancy positions were manually trimmed from motif 5′ ends. **e,** Schematic of *E. coli*-based plasmid interference assay using pEffector and pTarget (left), and bar graph plotting surviving colony-forming units (CFU) for the indicated conditions and proteins (right). TnpB nucleases cause robust cell death, whereas TldR homologs have no effect on cell viability, indicating a lack of DNA cleavage activity. EV, empty vector; M, TnpB mutant; NT, non-targeting guide; T, targeting guide. Bars indicate mean ± s.d. (n = 3). **f,** Alternative models of TldR-mediated transcriptional repression by blocking either transcription initiation or elongation by RNAP (blue). **g,** Schematic of RFP repression assay in which gRNAs were designed to target either the top or bottom strand of a promoter driving *rfp* expression (left), and bar graph plotting normalized RFP fluorescence for the indicated conditions. EV, empty vector; NT, non-targeting guide; Top/Btm, gRNA targeting the top or bottom strand. Bars indicate mean ± s.d. (n = 3). **h,** Experiments and data shown as in **g**, but with guides targeting the top/bottom strand within the 5′ UTR, downstream of the promoter. Results with nuclease-dead dCas12 and dCas9 are shown for comparison. Bars indicate mean ± s.d. (n = 3 for TldR; n = 6 for dCas12/dCas9).

Unexpectedly, RIP-seq revealed that the *oppF*-associated *Efa1*TldR bound an even shorter gRNA, comprising a 100-nt scaffold and ∼9-nt guide (**Extended Data Fig. 3a**); a similarly truncated guide (11 nt) was also observed for another homolog from this clade using publicly available RNA-seq data (**Extended Data Fig. 3b**). RIP-seq data from replicates and five additional homologs corroborated the short guide for *Efa1*TldR while revealing more heterogeneous processing for diverse homologs, including some with guides closer in length to 16-nt, others with more diffuse peaks that rendered unambiguous determination of the gRNA boundaries challenging, and one homolog (*Esa*TldR) that did not appear to specifically associate with its gRNA sequence (**Supplementary Data Fig. 3**).

Armed with the knowledge that *oppF*-associated TldRs associate with shortened guides, we performed a new search for putative genomic targets by screening for sites with ∼9-bp of DNA complementary to the guide flanked by a TAM similar to that recognized by related TnpB nucleases (TTTAA or TTTAT) (**Extended Data Fig. 4a**). This analysis led to the identification of a conserved target upstream of the start codon of one of the ABC transporter genes (*oppA*) encoded proximally to *tldR* (**Extended Data Fig. 4b,c**). OppA is a substrate binding protein (SBP) in ABC transport systems, and *tldR-*associated OppA homologs are most similar to SBPs that bind short polypeptides^39,48^ (**Extended Data Fig. 4d**). We hypothesized that TldR binding upstream of *oppA* could also serve a regulatory role, and when we expanded our analyses across diverse homologs, we found that the putative gRNA-matching targets varied in their orientation relative to the start codon of *oppA*, suggesting that TldRs from this clade might be able to target either DNA strand to transcriptionally repress *oppA*. Bioinformatic predictions with BPROM^49^ revealed that putative TldR targets indeed overlapped with the predicted –10 and –35 promoter elements of *oppA*, a conclusion corroborated by analysis of published RNA-seq data^46^ (**Extended Data Fig. 4e**). Interestingly, we also identified additional putative gRNA targets in genomes encoding *oppA-tldR* loci — including targets upstream of other ABC transporter components — raising the possibility that TldR proteins contribute towards a more complex transcriptional regulatory network than *fliC_P_*-associated TldR proteins (**Extended Data Fig. 5**).

Taken together, these data strongly support the hypothesis that TldR proteins across multiple independent lineages function as RNA-guided transcription factors to regulate gene expression, and that their biological targets relate to the accessory genes they stably associate with.

### TldRs function as RNA-guided DNA binding proteins that repress transcription

We selected seven *fliC_P_-*associated (**Fig. 2c**) and eight *oppF*-associated (**Extended Data Fig. 1a**) TldR homologs for functional assays, which were chosen to sample the diversity within each clade (**Supplementary Data Fig. 4**), cloned them into expression vectors alongside their putative gRNAs, and expressed them in an *E. coli* K12 strain containing a genomically integrated target site. We profiled genome-wide binding specificity using chromatin immunoprecipitation sequencing (ChIP-seq), and the resulting data revealed strongly enriched peaks corresponding to the expected target site for nearly all homologs tested (**Fig. 4b, Supplementary Data Fig. 5**). These data demonstrate that TldR proteins retain the ability to perform highly specific, RNA-guided DNA target binding in cells, despite harboring RuvC mutations and C-terminal truncations.

We next analyzed prominent off-target peaks in the ChIP-seq dataset. Strikingly, one of these off-target peaks for *fliC_P_*-associated TldRs corresponded to the intergenic region between *E. coli fliC* and *fliD* (**Fig. 4b,c**). The guide sequence used in these experiments is complementary to the native *fliC* target from *Enterobacter cloacae* sp. AR_154 but mutated relative to the *E. coli* K12 sequence at five positions (**Fig. 4c**), suggesting a high tolerance for TldR binding to mismatched targets (**Supplementary Data Fig. 5**). Strongly enriched peaks corresponding to off-target binding for *oppF*-associated TldRs similarly exhibited sequence similarity across only the TAM-proximal region of the target site (**Extended Data Fig. 6**). These data support the definition of a ∼6-nt TldR seed sequence, consistent with the previously reported 6-nt seed for some Cas12a homologs^50^.

ChIP-seq also captures transient interactions due to the crosslinking step, and we reasoned that systematic analysis of all peaks could report on the underlying TAM specificity of select TldR homologs, as we previously showed for TnpB^3^. Using MEME to detect enriched motifs, we found that *fliC -*associated TldRs were enriched at 5′-GTTAT-3′ motifs, the same pentanucleotide TAM that flanks putative TldR-gRNA targets within *fliC* promoters (**Fig. 4d, Supplementary Data Fig. 5**). Similarly, *oppF*-associated TldR homologs bound DNA sequences enriched in 5′-TTTAA-3′ motifs, which is consistent with the bioinformatically predicted TAM specificities for their closely related TnpB relatives (TTTAA and TTTAT) (**Supplementary Data Fig. 6**). Considered together, these results indicate that TAM sequences for TldR proteins can be accurately predicted *in silico*, even without the transposon context clues used for TnpB nucleases^2,3^.

To verify that the naturally-occurring RuvC mutations in TldR proteins actually abolish nuclease activity, we tested TldR homologs or their related TnpB counterparts in plasmid interference assays. Expression vectors containing TldR or TnpB and their associated gRNA (pEffector) were used to transform *E. coli* cells, along with a target plasmid (pTarget) bearing a kanamycin resistance cassette (*kanR*) and a TAM-flanked target sequence (**Fig. 4e**). Nuclease activity is expected to eliminate pTarget, resulting in fewer surviving colonies when cells are plated on selective media. When cells were transformed with plasmids bearing a previously studied TnpB homolog^3^ (i.e., *Gst*TnpB3) or nuclease-active TnpB homologs similar to TldRs (i.e., *Eko*TnpB2 and *Ece*TnpB), we were unable to isolate any surviving colonies, and this effect could be reversed using non-targeting guides or empty vector controls (**Fig. 4e**). In contrast, cells transformed with plasmids encoding TldR homolog exhibited similar colony counts as empty vector controls, with or without a pTarget-matching gRNA (**Fig. 4e**). Thirteen additional TldR homologs yielded consistent results (**Extended Data Fig. 7**), confirming that TldR proteins function as RNA-guided DNA binding proteins that lost the ability to cleave DNA.

Finally, we set out to investigate whether target DNA binding by TldR could modulate gene expression. Based on the overlap between putative TldR-gRNA targets and the predicted promoters driving *fliC* and *oppA* expression, we hypothesized that they would natively repress target gene expression, akin to the engineered use of nuclease-dead Cas9/Cas12 variants in CRISPR interference (CRISPRi) applications^51,52^. To test this, we developed an RFP/GFP reporter assay, in which target DNA binding represses *rfp* gene expression relative to a control *gfp* locus, and designed gRNAs to either occlude transcription initiation by targeting promoter sequences, or to block transcription elongation by targeting the 5′-untranslated regions (UTR) (**Fig. 4f,g**). We found that representative *fliC_P_*- (*Eho*) and *oppF*-associated (*Efa1*) TldR homologs robustly repressed RFP fluorescence when targeting the top (i.e., sense) strand, whereas only *Efa1*TldR repressed RFP when targeting the bottom (i.e., antisense) strand (**Fig. 4g**), consistent with the observation that putative native targets of *oppF*-associated TldRs occur on either strand relative to *oppA*. When we instead targeted the 5′-UTR, select TldRs from both clades only efficiently repressed RFP when targeting the bottom strand, whereby the TAM-proximal end was oriented towards the promoter and elongating RNAP, at efficiencies that were comparable to dCas9 and dCas12 (**Fig. 4h, Extended Data Fig. 8**).

Collectively, our results demonstrate that TldRs lack any detectable cellular nuclease activity, and instead function as RNA-guided DNA binding proteins with the potential to potently repress gene expression, in a mechanism reminiscent of engineered, nuclease-dead CRISPR-Cas effectors.

### Prophage-encoded *tldR* genes selectively repress host *fliC* expression *in vivo*

Having analyzed TldRs in heterologous contexts, we next turned to investigating their native function, focusing specifically on *fliC*-*tldR* loci. FliC, or flagellin, is the major extracellular subunit that polymerizes in tens of thousands of copies to form mature flagellar filaments, enabling bacterial locomotion (**Fig. 5a**) ^8^. Previous structural studies defined four domains of FliC proteins^53,54^, with D0 and D1 forming the majority of inter-promoter contacts during FliC polymerization, and D2 and D3 forming the central region that is predominantly exposed to the external environment (**Fig. 5b**). Remarkably, when we compared host FliC and prophage FliC_P_ sequences, we found that D2-3 were highly variable whereas D0-1 were highly conserved (**Fig. 5b,c**), suggesting that prophage flagellin would likely retain the ability to form flagella together with host components, while nevertheless diversifying the chemical composition of exposed filament surfaces. Flagellin D2-3 variation has long been recognized as a potential mechanism to evade mammalian host immune systems, since FliC is a primary antigen (i.e., antigen H) decorating pathogenic bacteria^10,55^. Moreover, some bacteriophages — eponymously referred to as flagellotropic phages — specifically recognize FliC within the flagellum as a primary receptor during adsorption^9,56^, likely through interactions with D2-3. We therefore hypothesized that prophage-encoded TldR proteins would repress host *fliC* expression as a way to remodel cellular flagella with the homologous protein products of TldR-associated *fliC* genes, at this crucial interface between host-pathogen (bacteria-phage) and pathogen-host (bacteria-mammal).

**Figure 5.**
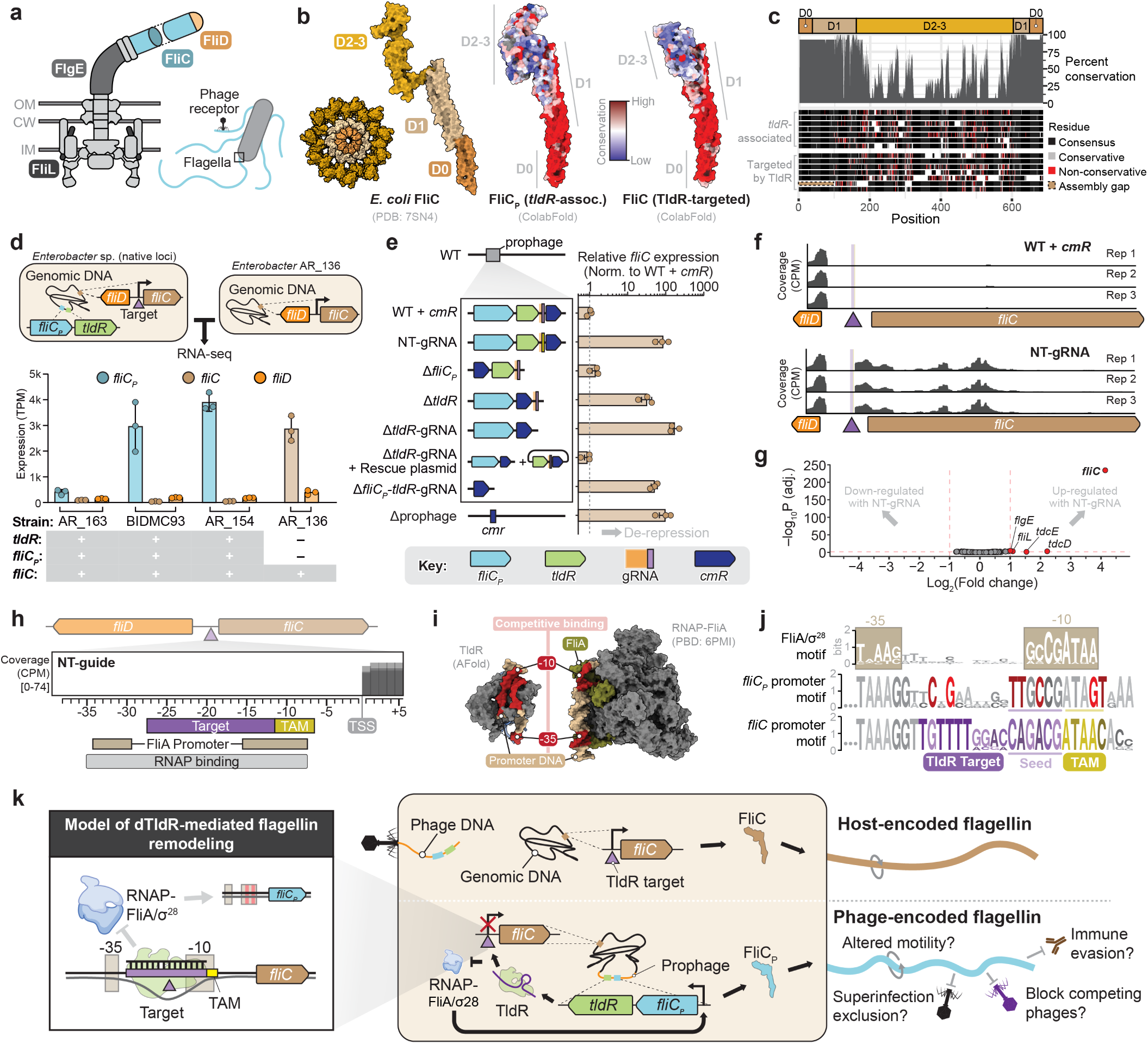
Flagellin-associated TldRs repress host flagellin gene expression in native clinical *Enterobacter* strains. **a,** Schematic of the flagellar assembly spanning the inner membrane (IM), cell wall (CW), and outer membrane (OM). The flagellin (FliC), hook (FlgE), stator-interacting (FliL), and flagellar cap (FliD) proteins are indicated. FliC filaments typically comprise several thousand subunits, are 5–20 µm in length, and are known receptors of flagellotropic phages. **b,** Surface representation of *E. coli* FliC (PDB: 7SN4) colored by domains, showing both a single monomer and filament cross section (left). Surface representations of ColabFold-predicted prophage FliC_P_ (middle) and host FliC (right) structures from *Enterobacter cloacae*, colored with AL2CO conservation scores calculated from the multiple sequence alignment (MSA) shown in **c**. **c,** MSA of TldR-associated FliC_P_ and TldR-targeted FliC proteins, showing the strongly conserved D0-1 domains and hypervariable D2-3 domains. **d,** Schematic of *Enterobacter* strains selected for RNA-seq analysis (top), and expression data plotted as transcripts per million (TPM) for *fliC_P_* (when present) and host *fliC* and *fliD*. The presence/absence of *fliC_P_*-*tldR* loci is indicated below the graph. Bars indicate mean ± s.d. (n = 3). **e,** Schematic of *Enterobacter cloacae* mutants generated by recombineering (left), and RT-qPCR analysis of host *fliC* expression levels normalized to the WT strain with *cmR* marker. Any deletion of *tldR* or substitution with a non-targeting (NT) gRNA leads to *fliC* de-repression. Bars indicate mean ± s.d. (n = 3). **f,** RNA-seq coverage at the host *fliC* locus for the indicated strains in **e**, showing de-depression with the NT-gRNA. **g,** Volcano plot showing differential gene expression analysis for the WT and NT-gRNA strains in **f**. Genes with a log_2_(fold change) ≥ 1 and an adjusted *p*-value < 0.05 are highlighted in red. **h,** Magnified view of data in **f**, showing the TAM/target overlap with predicted FliA/σ^28^ promoter elements inferred from *E. coli* K12 data. **i,** Predicted AlphaFold structure of TldR bound to target DNA (left) compared to experimental structure of RNAP (grey) and FliA/σ^28^ (green) bound to promoter DNA (right). **j,** Comparison of promoter motifs for host *fliC* and prophage *fliC_P_* alongside the FliA/σ^28^ motif from Tomtom analysis. This analysis suggests that *fliC_P_* is expressed similarly as *fliC*, while harboring conserved mutations (red) in the TAM and seed sequence that preclude self-targeting by its associated TldR. **k,** Model for the role of TldR in RNA-guided repression of host *fliC* upon temperate phage infection, leading to the selective expression and generation of phage-encoded flagellin (FliC_P_) filaments. The full range of biological implications from this transformation (right) will require further investigation.

We obtained and cultured three *Enterobacter* strains that each harbored a prophage-encoded *fliC_P_*-*tldR* locus, alongside a closely related control strain that lacked it, and performed total RNA-seq. Each strain with *tldR* exhibited robust gRNA expression, with 5′ and 3′ boundaries that were in excellent agreement with our heterologous RIP-seq data (**Extended Data Fig. 9**). Remarkably, when we analyzed flagellin gene expression relative to the flagellar hook (*fliD*), we found that host *fliC* was nearly undetectable in all three strains that encoded *tldR* whereas *fliC_P_* was strongly expressed (**Fig. 5d**), consistent with our hypothesis on TldR-gRNA function. In contrast, *fliC* was highly expressed in the control strain that lacked TldR and the prophage (**Fig. 5d**).

Next, to prove that *fliC* down-regulation was a direct consequence of TldR-mediated repression, rather than an indirect effect relating to the complex regulatory network controlling flagellar assembly^57^, we generated precise genetic perturbations to the *fliC_P_*-*tldR* locus in *Enterobacter cloacae* strain BID- MC93 and measured the corresponding effects on host *fliC* expression by RT-qPCR. Deletion of *tldR*, *tldR*-gRNA, the entire *fliC_P_*-*tldR-*gRNA locus, or the entire prophage, all led to a ∼100-fold increase in host *fliC* expression, and crucially, the same increase was observed after substituting the guide portion of the gRNA with a non-targeting (NT) control sequence (**Fig. 5e**). In contrast, deletion of *fliC_P_* alone had no effect, and the *fliC* expression increase could be reversed by complementing the *tldR-*gRNA deletion with a plasmid-encoded *tldR*-gRNA cassette (**Fig. 5e**). When we performed RNA-seq on isogenic strains that differed only in the guide sequence, across three biological replicates, we obtained clear evidence of host *fliC* de-repression with the NT-guide (**Fig. 5f**). Differential gene expression analyses further revealed that *fliC* was the most strongly up-regulated (i.e. de-repressed) gene transcriptome-wide (**Fig. 5g**), with the only other significant changes arising in genes whose expression has been linked to flagellar gene transcription^58,59^.

Closer inspection of the RNA-seq data lent further support to our conclusion that TldR represses gene expression through competitive binding to promoter elements, since the *fliC* transcription start site (TSS) agreed with the −35 and −10 promoter annotations informed from FliA/σ^28^ data in *E. coli* K12 (**Fig. 5h, Extended Data Fig. 10**). This interpretation was also corroborated by comparisons of predicted TldR-gRNA-DNA structures with an experimentally determined RNAP-FliA-DNA holoenzyme structure, which demonstrate that TldR target binding would sterically block FliA access to DNA (**Fig. 5i**). To determine how prophage-encoded *fliC_P_* genes would escape TldR-mediated repression, we applied MEME to detect conserved motifs in the region upstream of the experimentally-determined *fliC_P_* TSS, and then used Tomtom to compare these motifs to a database of known transcription factor binding sites. These analyses revealed that prophages likely recruit the very same host FliA/σ^28^ transcriptional program to produce FliC_P_, but with highly conserved mutations in both the TAM and seed sequence that preclude TldR-gRNA recognition (**Fig. 5j**). Collectively, then, *fliC_P_-tldR* locus is elegantly adapted to remodel composition of the flagellar apparatus upon establishment of a lysogen, by selectively repressing host flagellin through RNA-guided DNA targeting while hijacking cellular machinery to express its own homolog substitute (**Fig. 5k**).

## DISCUSSION

Bacterial flagella represent a critical nexus at the host-pathogen interface, and the attendant selective pressures likely contributed to the domestication and emergence of *fliC-*associated *tldR* genes on at least two independent occasions, to sensitively regulate flagellar expression and composition (**Fig. 2a**). A number of bacteriophages, including the well-studied *Salmonella* Chi phage, recognize the flagellar filament as a receptor during cell absorption^9,56,60^, and phage-mediated substitution of host flagellin may thus prevent superinfection and/or render the cell invisible to competing flagellotropic phages in the environment (**Fig. 5k**). Flagellin/FliC, known as the H antigen in bacterial pathogen serotyping, also functions as a primary antigen that is recognized by both receptors and antibodies in the mammalian innate and adaptive immune systems^61–64^, and the pervasive presence of prophage-encoded *fliC_P_-tldR* loci in clinical isolates from humans (**Supplementary Dataset 2**) could represent a novel example of lysogenic conversion^65^, whereby phages enable their bacterial hosts to evade an immune response. Finally, flagellar remodeling could also modulate motility of bacterial host cells, and thus impact their capabilities for chemotaxis and nutrient acquisition. Resolving how RNA-guided repression of host flagellin gene expression impacts one or more facets of bacterial physiology, and whether flagellar composition is dynamically altered over time in these lysogenic strains, will be a major goal of future research efforts (**Fig. 5k**).

The biological purpose of *oppF*-associated TldR homologs that are encoded next to ABC transporter operons (**Fig. 2a**) is less clear, though our gRNA discovery approach — blending both bioinformatics predictions and experimental validation via RIP-seq — revealed a likely function in controlling expression of the key periplasmic binding protein, OppA (**Extended Data Fig. 4**). ABC transporters are ubiquitous membrane-bound protein complexes that move substrates in and out of cells, and are responsible for nutrient uptake, drug resistance, toxin efflux, and virulence factor secretion, among many other roles^39,66^. TldR proteins may provide a mechanism to regulate *oppA* expression in response to external biological cues, though more experiments will be needed to better understand their activities and specificities *in vivo*, especially given the truncated guide sequences they use. More generally, the advantages offered by RNA-guided transcription factors that seem to primarily function with just a single gRNA, as compared to more canonical sequence-specific protein transcription factors, will require additional investigation.

Our work reveals that transposon-encoded TnpB nucleases have been repurposed multiple times for new functions in gene regulation, and that nuclease-inactivating mutations in the RuvC domain coincided with novel gene associations. By integrating knowledge about the biochemical properties of closely related TnpB nucleases, including their TAM and ωRNA requirements, together with systematic biochemical profiling using ChIP-seq and reporter assays, we were able to straightforwardly identify the gRNAs and targets recognized by TnpB-like nuclease-dead repressor (TldR) proteins, providing an important advance beyond descriptive bioinformatic observations^30^. Together, these studies provide an expansive view on the biological opportunities afforded by RNA-guided DNA targeting, from ensuring the proliferation of transposons in their primordial context, to protecting cells against foreign nucleic acids in their CRISPR-Cas context, to selectively modulating gene expression in their nuclease-dead TldR context. It appears certain that additional examples of TnpB domestication will be uncovered with further bioinformatic and experimental mining, for both bacterial and eukaryotic Fanzor homologs^27,28^, and future efforts should be broadened to include nuclease-dead and nuclease-active variants that exist in non-transposon and non-mobile genetic contexts.

It is noteworthy that evolution has repeatedly sampled some of the very same molecular innovations invented by humans during the development of CRISPR-based genome engineering technologies. A decade ago, Qi and colleagues developed the first applications of synthetic, nuclease-dead variants of Cas9 (i.e. dCas9) for transcriptional modulation^51^, and intense efforts ever since have resulted in a plethora of highly effective tools for epigenome editing, typically via engineered fusions of dCas9 to diverse effector domains^67^. In its native bacterial context, though, Cas9 already exploits a mechanism of autoregulatory gene expression control using natural long-form tracrRNAs with truncated guides to bind, but not cleave, its own *cas9* promoter sequence^68^. Other non-canonical guide RNAs similarly program Cas9 for natural gene repression functions as a means of promoting virulence^69^, and some CRISPR-Cas subtypes leverage nuclease-dead Cas12 variants for adaptive immune protection, in a mechanism that relies on high-affinity RNA-guided DNA binding without cleavage^32,33^. The recent observation of *cas12f*/*tnpB*-like genes adjacent to sigma factor genes^30^ suggests the exciting possibility that nuclease-dead, RNA-guided DNA targeting proteins have also already been sampled for gene activation. Taken together with our discoveries of TldR function, these examples reveal that transcriptional down-regulation and up-regulation via programmable CRISPRi and CRISPRa approaches, respectively^35^, emerged in nature long before humans deciphered the molecular mechanisms of CRISPR-Cas.

## METHODS

### Bioinformatic identification of natural, nuclease-dead TnpB homologs (TldRs)

An initial search of the NCBI non-redundant (NR) protein database — queried with TnpB sequences from *H. pylori* and *G. stearothermophilus* (WP_078217163.1 and WP_047817673.1, respectively) in Jackhmmer^70^ — resulted in the identification of 95,731 unique TnpB-like proteins, which were further clustered at 50% amino acid identity (across 50% sequence coverage) via CD-HIT^71^ to produce a set of 2,646 representative TnpB sequences. A multiple sequence alignment (MSA) was then constructed with MAFFT^72^ (EINSI; four rounds), which was trimmed manually with trimAl^73^ (90% gap threshold; v1.4.rev15). The resulting alignment of TnpB/TldR homologs was used to construct a phylogenetic tree in IQTree^74^ (WAG model, 1000 replicates for SH-aLRT, aBayes, and ultrafast bootstrap) ^3^, which was annotated and visualized in ITOL^75^.

To assess the conservation of RuvC catalytic residues in each TnpB protein sequence, we compared each sequence in the MSA to structurally characterized orthologs (i.e., *Dra*TnpB from ISDra2 and Cas12f; PDB ID 8H1J and 7L48, respectively). This comparison was performed by aligning each candidate, as well as the homologs represented in the closest five tree branches on either side of it, to *Dra*TnpB and *Un*Cas12f using the AlignSeqs function of the DECIPHER package^76^ in R. TnpB-like protein sequences with less than two conserved residues of the RuvC DED catalytic motif were extracted using the Biostrings package^77^ in R. For each sequence with less than two active site residues identified (defined as a TnpB-like nuclease-dead Repressor, or TldR), related homologs were retrieved from initial sequence clusters, and additional related homologs were identified via BLASTP searches of the NR protein database (e-value < 1e-50, query coverage > 80%, max target sequences = 50) ^78^. Each representative sequence and all of their cluster members were used as queries in these BLASTP searches, and the active sites from BLAST hits were checked by aligning proteins to structurally determined representatives, as described above. This approach resulted in the identification of 366 unique TldR homologs. Genomes encoding each TldR were retrieved from NCBI using the batch-entrez tool. TldR-encoding loci (i.e., *tldR* +/- 20 kbp) were extracted using the Biostrings package^77^ in R, and each *tldR* locus was annotated with Eggnog (-m diamond --evalue 0.001 --score 60 --pident 40 --query_cover 20 --subject_cover 20 --genepred prodigal --go_evidence non-electronic --pfam_realign none) ^79^. Annotated *tldR* loci were manually inspected in Geneious.

### Bioinformatic analyses of *fliC_P_*-, *oppF*-, and *csrA-*associated TldR homologs

To further investigate *fliC*-associated TldR homologs, we extracted cluster members for three representative branches in the tree shown in **Figure 1** (WP_193971683.1, WP_064735610.1, and WP_048785942.1). The protein file representing these combined clusters was supplemented with additional homologs identified via BLASTP searches of the NR database^78^. The resulting concatenated protein file included both TldR and related TnpB sequences. To increase the diversity of TnpB proteins represented in this dataset, three additional TnpB homologs (WP_269608765.1, WP_024186316.1, WP_059759460.1) were identified and manually added to this protein file via web-based BLASTP searches queried with the TnpB protein sequences already present in the dataset (e-value < 0.05). An MSA was constructed from these sequences and *Dra*TnpB using the AlignSeqs function of the DECIPHER package^76^ in R to verify the active site composition of each ortholog. To determine which *tldR/tnpB* genes were associated with *fliC*, we analyzed Eggnog annotation information for each locus (described above) and extracted TldR/TnpB sequences that were encoded within three open reading frames of *fliC*.

A locus was defined as phage-associated if it contained four or more gene annotations that contained the word “Phage”, “phage”, “Viridae”, or “viridae”. TldR/TnpB protein sequences were then de-duplicated via CD-HIT^71^ (-c 1.0), and an MSA was built in MAFFT^72^ (LINSI) from the resulting set of 160 unique proteins. Protein domain coordinates displayed around the tree in **Figure 2c** were inferred by cross-referencing the MSA and predicted structures. The phylogenetic tree shown in **Figure 2c** was built from the TldR/TnpB MSA in FastTree^80^ (-wag -gamma) and was annotated and visualized in ITOL^75^. Structural models of each candidate shown in **Figure 1d** were predicted with AlphaFold^81^ (v2.3) and displayed with ChimeraX^82^ (v1.6); MSAs were visualized in Jalview^83^.

To interrogate *oppF*-associated TldR sequences, we extracted cluster members and additional homologs identified via BLASTP^78^ searches of the NR database (e-value < 1e-50, query coverage > 80%, max target sequences = 50) for six branches representing TldR proteins in the **Figure 1c** tree (RBR34854.1, WP_016173224.1, WP_156233666.1, NTQ19983.1, OTP13636.1, OSH30650.1). We concatenated these sequences with cluster members and additional homologs identified through an identical BLASTP search of one representative TnpB branch (EOH94253.1) that corresponded to the closest branch to the six TldR branches in the tree. To increase the diversity of related TnpB proteins represented in this dataset, three additional TnpB homologs (WP_242450195.1, WP_028983493.1, WP_277281207.1) were identified and manually added to this protein file via web-based BLASTP searches queried with the TnpB protein sequences already present in the dataset (e-value < 0.05). Genomes encoding TldR/TnpB proteins were downloaded from NCBI using the Batch-entrez tool, relevant loci (*tldR/tnpB* +/- 20 kbp) were extracted using the Biostrings package^77^ in R, and each locus was annotated with Eggnog (see above) ^79^. Each TldR/TnpB protein was individually aligned to *Dra*TnpB using the AlignSeqs function of the DECIPHER package^76^ in R to verify its RuvC active site composition. TldR/TnpB sequences were then deduplicated via CD-HIT^71^ (-c 1.0), and an MSA was built in MAFFT^72^ (LINSI) from the resulting set of 204 unique proteins. An initial phylogenetic tree was constructed in FastTree^80^ (-wag -gamma), and this tree was used to guide the selection of eight representative TldRs and four representative TnpBs (shown in **Supplementary** Figure 4) that were structurally predicted with ColabFold^84^ (v1.5). These twelve predicted structures were used to guide an alignment of TldR/ TnpB protein sequences in Promals3D^85^, and the resulting MSA was used to build the tree in **Extended Data Figure 1** in FastTree (-wag -gamma). Protein domain coordinates displayed around the tree in **Extended Data Figure 1** were inferred by cross referencing the MSA and predicted structures. The phylogenetic tree was annotated and visualized in ITOL^75^.

To probe *oppF*-associated TldR loci, we extracted cluster members and additional homologs identified via BLASTP^78^ searches of the NR database (e-value < 1e-50, query coverage > 80%, max target sequences = 500) for one TldR protein in the **Figure 1c** tree (WP_204886977.1). Genomes encoding TldR/TnpB proteins were downloaded from NCBI using the Batch-entrez tool, relevant loci (*tldR/ tnpB* +/- 20 kbp) were extracted using the Biostrings package^77^ in R, and each locus was annotated with Eggnog (see above) ^79^. Each TldR/TnpB protein was individually aligned to *Dra*TnpB using the AlignSeqs function of the DECIPHER package^76^ in R to verify its RuvC active site composition. TldR/TnpB sequences were then deduplicated via CD-HIT^71^ (-c 1.0), resulting in 36 unique additional TldR proteins.

### Bioinformatic identification of TldR-associated gRNA sequences

To define the boundaries of gRNA scaffolds in *fliC_P_-tldR* loci, we used a general gRNA covariance model (CM) described in previous work^3^. The CMsearch function of Infernal (Inference of RNA alignments; v1.1.2) ^86^ was used to scan nucleotide sequences of *tldR* and 500-bp flanking windows, resulting in the identification of putative gRNA scaffold sequences. These TldR-associated gRNA scaffold boundaries were confirmed by comparing *fliC_P_*-*tldR* loci to ωRNAs from confidently predicted annotations of catalytically active TnpB loci. Putative TldR guide sequences could then be retrieved from the 3′ boundary of putative gRNA scaffolds, enabling prediction of native *fliC_P_-*associated TldR targets. Putative guides are listed in **Supplementary Dataset 2**).

An analogous search of *oppF*-associated *tldR* loci with a general gRNA CM failed to identify putative gRNA sequences. For this group of *tldR* loci, we instead built a new CM from ωRNA sequences associated more closely related TnpB loci. Using the comparative genomics strategy outlined in **Figure 3a**, we manually identified the putative transposon right end (RE) for one TnpB-encoding IS element (WP_113785139.1 in KZ845747). We then aligned nucleotide sequences for all the related *tnpB* genes and 500 bp of sequence downstream of *tldR* with MAFFT^72^ (LINSI). The resulting alignment was trimmed at the 3′ end to the position of the ωRNA scaffold-guide boundary identified for the WP_113785139.1 locus. This putative set of TnpB ωRNA sequences was used realigned with LocaRNA^87^ (--max-diff-at- am=25 --max-diff=60 --min-prob=0.01 --indel=-50 --indel-opening=-750 --plfold-span=100 --alifold-consensus-dp; v2.0.0), and a CM (ABC_gRNA_v1) was built and calibrated with Infernal. The CMsearch function of Infernal was then used to search sequences composed of *tldR/tnpB* and 500 bp of down-

stream sequence with the ABC_gRNA_v1 CM. This search resulted in gRNA identification for some, but not all, *tldR* loci. Thus, a second gRNA CM was built by extracting the newly identified TldR/TnpB gRNA sequences from their respective genomes, merging them with the sequences used to construct ABC_gRNA_v1, aligning the prospective gRNA dataset in LocaRNA, and building and calibrating a new CM with Infernal (ABC_gRNA_v2). When sequences comprising *tldR/tnpB* and 500 bp downstream were scanned with the ABC_gRNA_v2 CM, via CMsearch, putative gRNA sequences were identified for the remaining *tldR* loci (listed in **Supplementary Dataset 3**).

### Visualization of RNA-seq data from the NCBI short read archive (SRA) and gene expression omnibus (GEO)

To assess gRNA expression from a representative *fliC_P_-tldR* locus, an RNA-seq dataset was downloaded from the NCBI SRA (accession: ERR6044061). Reads were aligned to the *Enterobacter cloacae* AR_154 genome (CP029716.1) with using bwa-mem2^88^ (v2.2.1) in paired-end mode with default parameters, and alignments were converted to BAM files with SAMtools^89^. Bigwig files were generated with the bamCoverage utility in deepTools^90^, and unique reads mapping to the forward strand were visualized with the Integrated Genome Viewer (IGV) ^91^. Expression of gRNA and *oppA* from an *oppF*-*tldR* locus was assessed by downloading an RNA-seq analysis from the NCBI GEO (accession: GSE115009). Normalized coverage files (ID-005241, ID-005244, ID-005245, ID-005246) for the forward strand were visualized in IGV^91^.

### Plasmid and *E. coli* strain construction

All strains and plasmids used in this study are described in **Supplementary Tables 1 and 2**, respectively, and a subset is available from Addgene. In brief, genes encoding candidate TldR and TnpB homologs (**Supplementary Table 3**), alongside their putative gRNAs, were synthesized by GenScript and subcloned into the PfoI and Bsu36i restriction sites of pCDF- Duet-1, to generate pEffector, similar to our previous work^3^. Expression vectors contained constitutive J23105 and J23119 promoters driving expression of *tldR*/*tnpB* and the gRNA, respectively, and *tldR*/ *tnpB* genes encoded an appended 3×FLAG-tag at the N-terminus. gRNAs for *fliC_P_*-associated TldRs were designed to target the host *fliC* 5′ UTR site, whereas gRNAs of *oppF*-associated TldRs were engineered to target the genomic site natively targeted by a *Gst*TnpB3 homolog. Derivatives of these pEffector plasmids, or their associated pTarget plasmids (for plasmid interference assays), were cloned using a combination of methods, including Gibson assembly, restriction digestion-ligation, ligation of hybridized oligonucleotides, and around-the-horn PCR. Plasmids were cloned, propagated in NEB Turbo cells (NEB), purified using Miniprep Kits (Qiagen), and verified by Sanger sequencing (GENEWIZ).

A custom *E. coli* K12 MG1655 strain that contained genomically-encoded *sfGFP* and *mRFP* genes was constructed by adding three target sites adjacent to bioinformatically predicted TAM sequences upstream of the *mRFP* ORF, in between the constitutive promoter driving RFP expression and the corresponding ribosome binding site (sSL3580; derivative of GenBank: NC_000913.3) ^51^ (**Supplementary Table 1**). The original strain (with genomic *sfGFP* and *mRFP*) was a gift from L. S. Qi. The inserted target sites represent 25-bp sequences derived from the 5′ UTR of host *fliC* (*Enterobacter cloacae* complex sp. strain AR_0154; GenBank: CP029716.1), an ABC transporter gene (*Enterococcus faecium* strain BP657; GenBank: CP059816.1), and a *Gst*TnpB3 native target used previously^3^.

### Chromatin immunoprecipitation sequencing (ChIP-seq) and motif analyses of genomic sites bound by TldR

ChIP-seq experiments and data analyses were generally performed as described previously^3,92^, except for the use of sSL3580. In brief, *E. coli* MG1655 cells were transformed with pEffector and incubated for 16 h at 37 °C on LB-agar plates with antibiotic (200 µg ml^−1^ spectinomycin). Cells were scraped and resuspended in LB broth. The OD600 was measured, and approximately 4.0 × 10^8^ cells (equivalent to 1 ml with an OD_600_ of 0.25) were spread onto two LB-agar plates containing antibiotic (200 µg ml^−1^ spectinomycin). Plates were incubated at 37 °C for 24 h. All cell material from both plates was then scraped and transferred to a 50-ml conical tube. Cross-linking was performed in LB medium using formaldehyde (37% solution; Thermo Fisher Scientific) and was quenched using glycine, followed by two washes in TBS buffer (20 mM Tris-HCl pH 7.5, 0.15 M NaCl). Cells were pelleted and flash-frozen using liquid nitrogen and stored at −80 °C.

Chromatin immunoprecipitation of FLAG-tagged TnpB and TldR proteins was performed using Dynabeads Protein G (Thermo Fisher Scientific) slurry (hereafter, beads or magnetic beads) conjugated to ANTI-FLAG M2 antibodies produced in mouse (Sigma-Aldrich). Samples were sonicated on a M220 Focused-ultrasonicator (Covaris) with the following SonoLab 7.2 settings: minimum temperature, 4 °C; set point, 6 °C; maximum temperature, 8 °C; peak power, 75.0; duty factor, 10; cycles/bursts, 200; 17.5 min sonication time. After sonication, a non-immunoprecipitated input control sample was frozen. The remainder of the cleared sonication lysate was incubated overnight with anti-FLAG-conjugated magnetic beads. The next day, beads were washed, and protein-DNA complexes were eluted. The non-immunoprecipitated input samples were thawed, and both immunoprecipitated and non-immunoprecipitated controls were incubated at 65 °C overnight to reverse-crosslink proteins and DNA. The next day, samples were treated with RNase A (Thermo Fisher Scientific) followed by Proteinase K (Thermo Fisher Scientific) and purified using QIAquick spin columns (QIAGEN).

ChIP-seq Illumina libraries were prepared for immunoprecipitated and input samples using the NEBNext Ultra II DNA Library Prep Kit for Illumina (NEB). Following adapter ligation, Illumina barcodes were added by PCR amplification (12 cycles). ∼450-bp DNA fragments were selected using two-sided AMPure XP bead (Beckman Coulter) size selection. DNA concentrations were determined using the DeNovix dsDNA Ultra High Sensitivity Kit and dsDNA High Sensitivity Kit. Illumina libraries were sequenced in paired-end mode on the Illumina NextSeq platform, with automated demultiplexing and adapter trimming (Illumina). >2,000,000 raw reads, including genomic- and plasmid-mapping reads, were obtained for each ChIP-seq sample.

Following sequencing, paired-end reads were trimmed and mapped to a custom *E. coli* K12 MG1655 reference genome (derivative of GenBank: NC_000913.3). Genomic *lacZ* and *lacI* regions partially identical to plasmid-encoded genes were masked in all alignments (genomic coordinates: 366,386-367,588). Mapped reads were sorted and indexed, and multi-mapping reads were excluded. Alignments were normalized by counts per million (CPM) and converted to 1-bp-bin bigwig files using the deepTools2^90^ command bamCoverage, with the following parameters: --normalizeUsing CPM -bs 1. CPM-normalized reads were visualized in IGV^91^. Genome-wide views were generated using plots of maximum read coverage values in 1-kb bins. Peak calling was performed using MACS3^93^ (version 3.0.0a7) using the non-immunoprecipitated control sample of *Eco*TldR as reference. 200-bp sequences for each peak were extracted from the *E. coli* reference genome using BEDTools^94^ (v2.30.0), and sequence motifs were identified using MEME-ChIP^95^ (5.4.1). Primers used for Illumina library preparation are listed in **Supplementary Table 4**, and ChIP-seq read and meta information is listed in **Supplementary Table 5.**

### RNA immunoprecipitation sequencing (RIP-seq) of RNA bound by TldR

Cells harvested for RIP-seq were cultured as described for ChIP-seq using an *E. coli* K12 MG1655 strain expressing *sfGFP* and *mRFP* (sSL3580). Colonies from a single plate were scraped and resuspended in 1 ml of TBS buffer (20 mM Tris-HCl pH 7.5, 0.15 M NaCl). Next, the OD_600_ was measured for a 1:20 mixture of the cell suspension and TBS buffer, and a standardized amount of cell material equivalent to 20 ml of OD_600_ = 0.5 was aliquoted. Cells were pelleted by centrifugation at 4,000 *g* and 4 °C for 5 min. The supernatant was discarded, and pellets were stored at −80 °C.

Antibodies for immunoprecipitation were conjugated to magnetic beads as follows: for each sample, 60 μl Dynabeads Protein G (Thermo Fisher Scientific) were washed 3× in 1 ml RIP lysis buffer (20 mM Tris-HCl pH 7.5, 150 mM KCl, 1 mM MgCl_2_, 0.2% Triton X-100), resuspended in 1 ml RIP lysis buffer, and combined with 20 μl anti-FLAG M2 antibody (Sigma-Aldrich), and rotated for >3 h at 4 °C. Antibody-bead complexes were washed 3× to remove unconjugated antibodies, and resuspended in 60 μl RIP lysis buffer per sample.

Flash-frozen cell pellets were resuspended in 1.2 ml RIP lysis buffer supplemented with cOmplete Protease Inhibitor Cocktail (Roche) and SUPERase•In RNase Inhibitor (Thermo Fisher Scientific). Cells were then sonicated for 1.5 min total (2 sec ON, 5 sec OFF) at 20% amplitude. Lysates were centrifuged for 15 min at 4 °C at 21,000 *g* to pellet cell debris and insoluble material, and the supernatant was transferred to a new tube. At this point, a small volume of each sample (24 μl, or 2%) was set aside as the “input” starting material and stored at −80 °C.

For immunoprecipitation, each sample was combined with 60 μl antibody-bead complex and rotated overnight at 4 °C. Next, each sample was washed 3× with ice-cold RIP wash buffer (20 mM Tris-HCl, 150 mM KCl, 1 mM MgCl_2_). After the last wash, beads were resuspended in 1 ml TRIzol (Thermo Fisher Scientific) and RNA was eluted from the beads by incubating at RT for 5 min. A magnetic rack was used to separate beads from the supernatant, which was transferred to a new tube and combined with 200 μl chloroform. Each sample was mixed vigorously by inversion, incubated at RT for 3 min, and centrifuged for 15 min at 4 °C at 12,000 *g*. RNA was isolated from the upper aqueous phase using the RNA Clean & Concentrator-5 kit (Zymo Research). RNA from input samples was isolated in the same manner using TRIzol and column purification. High-throughput sequencing library preparation was performed as described below for total RNA-seq of *Enterobacter* strains. Libraries were sequenced on an Illumina NextSeq 550 in paired-end mode with 75 cycles per end.

Adapter trimming, quality trimming, and read length filtering of RIP-seq reads was performed as described below for total RNA-seq experiments. Trimmed and filtered reads were mapped to a reference containing both the MG1655 genome (NC_000913.3) and plasmid sequences using bwa-mem2 v2.2.1, with default parameters. Mapped reads were sorted, indexed, and converted into coverage tracks as described below for total RNA-seq experiments.

### Plasmid cleavage assays

Plasmid interference assays were generally performed as previously described^3^. *E. coli* K12 MG1655 (sSL0810) cells were transformed with pTarget plasmids (vector sequences are listed in **Supplementary Table 2**), and single colony isolates were selected to prepare chemically competent cells. Next, cells were transformed with 400 ng of pEffector plasmid or empty vector. After 3 h recovery at 37 °C, cells were pelleted by centrifugation at 4,000 *g* for 5 min and resuspended in 100 µl of H_2_O. Cells were then serially diluted (10×), plated as 8 µl spots onto LB agar supplemented with spectinomycin (200 µg ml^−1^) and kanamycin (50 µg ml^−1^), and grown for 16 h at 37 °C. Plate images were taken using a BioRad Gel Doc XR+ imager.

Plasmid interference assays were quantified by determining the number of colony-forming units (CFU) following transformation. Experiments were performed as described above, however for each experiment, 30 µl of a 10-fold dilution were plated onto a full LB agar plate containing spectinomycin (200 µg ml^−1^) and kanamycin (50 µg ml^−1^). CFUs were counted following 16 h of growth at 37 °C and reported as CFUs per µg of transformed pEffector plasmid.

### RFP repression assays

The RFP repression assay protocol was adapted from our previous study^3,92^. An *E. coli* strain expressing a genomically-integrated *sfGFP* (sSL3761), derived from a strain kindly provided by L. S. Qi^51^, was co-transformed with 200 ng of pEffector and pTarget (vector sequences listed in **Supplementary Table 2**). Protein components and guide RNAs (gRNA, sgRNA or crRNA) were constitutively expressed from pEffector. pTargets were cloned to encode an *mRFP* gene under the control of a constitutive promoter. For RFP repression assays shown in **Figure 4g**, gRNAs were designed to target the constitutive RFP promoter on either strand, and 5-bp TAM sequences were inserted 5′ of each target site. For RFP repression assays shown in **Figure 4h**, 25-bp sequences containing the TAM/ PAM and target site in either orientation were inserted in between the *mRFP* promoter and ribosome binding site.

Transformed cells were plated on LB-agar with antibiotic selection, and at least three of the resulting colonies on each plate were used to inoculate overnight liquid cultures. For each sample, 1 µl of the overnight culture was used to inoculate 200 µl of LB medium on a 96-well optical-bottom plate. The fluorescence signals for sfGFP and mRFP were measured alongside the OD_600_ using a Synergy Neo2 microplate reader (Biotek), while shaking at 37 °C for 16 h. For all samples, the fluorescence intensities at OD_600_ = 1.0 were used to determine the fold repression for each TldR or Cas targeting complex, and the data were normalized to the non-repressed signal for sSL3761. Background GFP and RFP fluorescence intensities at OD_600_ = 1.0 were determined using an *E. coli* K12 MG1655 strain (sSL0810) lacking *sfGFP* and *mRFP* genes, and were subtracted from all RFP and GFP fluorescence measurements.

### Total RNA sequencing of *Enterobacter* strains

*Enterobacter cloacae* strains (sSL3710, sSL3711, and sSL3712) were obtained from a CDC isolate panel (*Enterobacterales* Carbapenemase Diversity; CRE in ARIsolateBank), and an *Enterobacter* sp. BIDMC93 (sSL3690) was kindly provided by Ashlee M. Earl at the Broad Institute; strain information is listed in **Supplementary Table 1**. Biological replicates were obtained by isolating 3 individual clones of each *Enterobacter* strain on LB-agar plates and using these to inoculate overnight cultures in liquid LB media. All strains were grown at 37 °C without antibiotics and with agitation when in liquid medium (240 rpm), in a BSL-2 environment. For total RNA-seq library preparation, RNA was purified from 2 mL of exponentially growing cultures of sSL3690, sSL3710, sSL3711, and sSL3712, since RT-qPCR analyses of *fliC* expression showed that the TldR-mediated was more robust in exponential than in stationary phase. RNA was extracted using TRIzol and column purification (NEB Monarch RNA cleanup kit), and samples were then individually diluted in NEBuffer 2 (NEB) and fragmented by incubating at 92 °C for 1.5 min. The fragmented RNA was simultaneously treated with RppH (NEB) and TURBO DNase (Thermo Fisher Scientific) in the presence of SUPERase•In RNase Inhibitor (Thermo Fisher Scientific), in order to remove DNA and 5′ pyrophosphate. For further end repair to enable downstream adapter ligation, the RNA was treated with T4 PNK (NEB) in 1× T4 DNA ligase buffer (NEB). Samples were column-purified using RNA Clean & Concentrator-5 (Zymo Research), and the concentration was determined using the DeNovix RNA Assay (DeNovix). Illumina adapter ligation and cDNA synthesis were performed using the NEBNext Small RNA Library Prep kit, using 100 ng of RNA per sample. High-throughput sequencing was performed on an Illumina NextSeq 550 in paired-end mode with 75 cycles per end.

RNA-seq reads were processed using cutadapt^96^ (v4.2) to remove adapter sequences, trim low-quality ends from reads, and exclude reads shorter than 15 bp. Trimmed and filtered reads were aligned to reference genomes (accessions listed in **Supplementary Table 1**) using bwa-mem2^88^ (v2.2.1) in paired-end mode with default parameters. SAMtools^89^ (v1.17) was used to filter for uniquely mapping reads using a MAPQ score threshold of 1, and to sort and index the unique reads. Coverage tracks were generated using bamCoverage^90^ (v3.5.1) with a bin size of 1, read extension to fragment size, and normalization by counts per million mapped reads (CPM) with exact scaling. Coverage tracks were visualized using IGV^91^. For transcript-level quantification, the number of read pairs mapping to annotated transcripts was determined using featureCounts^97^ (v2.0.2). The resulting counts values were converted to transcripts-per-million-mapped-reads (TPM) by normalizing for transcript length and sequencing depth. For differential expression analysis between genetically engineered *Enterobacter* strains, the counts matrix was first filtered to remove rows with fewer than 10 reads for at least 3 samples. The filtered matrix was then processed by DESeq2^98^ (v1.40.2) in order to determine the log_2_ (fold change) for each transcript between the experimental conditions, as well as the Wald test *P* value adjusted for multiple comparisons using the Benjamini-Hochberg approach. Significantly differentially expressed genes were determined by applying thresholds of |log_2_(fold change)| > 1 and adjusted *P* value < 0.05.

### Construction of *Enterobacter* BIDMC93 mutants

*Enterobacter cloacae* strains AR_154 and AR_163 (sSL3711 and sSL3712; respectively) are both resistant to the antibiotics commonly used for colony selection following plasmid transformation, so we proceeded with recombineering in *Enterobacter* sp. BIDMC93. Genomic mutants (listed in **Supplementary Table 1**) were generated using Lambda Red recombineering, as previously described^99^. Mutants were designed to introduce a chloramphenicol resistance cassette at each disrupted locus. The chloramphenicol resistance cassette was amplified by PCR with Q5 High Fidelity DNA Polymerase (NEB), using primers that contained at least 50-bp of homology to the disrupted locus. Amplified products were resolved on a 1% agarose gel and purified by gel extraction (QIAGEN). Electrocompetent *Enterobacter* sp. BIDMC93 cells were prepared containing a temperature-sensitive plasmid encoding Lambda Red components under a temperature-sensitive promoter (pSIM6). Immediately prior to preparing electrocompetent cells, Lambda Red protein expression was induced by incubating cells at 42 °C for 25 min. 200-500 ng of each insert was used to transform cells via electroporation (2 kV, 200 Ω, 25 µF). Cells were recovered by shaking in 1 mL of LB media at 37 °C overnight. After recovery, cells were spread on 100 mm plates with 25 µg/mL chloramphenicol and grown at 37 °C. Chloramphenicol-resistant colonies were genotyped by Sanger sequencing (GE-NEWIZ) to confirm the desired genomic mutation.

### RT-qPCR to assess host *fliC* transcription in *Enterobacter* sp. BIDMC93

200 ng of the purified total RNA was used as an input for the reverse transcription reaction. First, total RNA was treated with 1 µl dsDNase (Thermo Fisher Scientific) in 1X dsDNase reaction buffer in a final volume of 10 µl and incubated at 37 °C for 20 min. Then, 1 µl of 10 mM dNTP, µl of 2 µM oSL14254, and 1 µl of 2 µM oSL14280 were added for gene-specific priming (*rrsA* and *fliC*, respectively), and reactions were heated at 65 °C for 5 min; oligonucleotide sequences are listed in **Supplementary Table 4**. Reactions were then placed directly on ice, followed by addition of 4 µl of SSIV buffer, 1 µl 100 mM DTT, 1 µl SUPERase•In™ (Thermo Fisher Scientific), and 1 µl of SuperScript IV Reverse Transcriptase (200 U/ µl, Thermo Fisher Scientific), followed by incubation at 53 °C for 10 min, and then incubation at 80 °C for 10 min. Quantitative PCR was performed in 10 µl reaction containing 5 µl SsoAdvanced™ Universal SYBR Green Supermix (BioRad), 1 µl H_2_0, 2 µl of primer pair at 2.5 µM concentration, and 2 µl of 100-fold diluted RT product. Two primer pairs were used: oSL14254/oSL14255 was used to amplify *rrsA* cDNA, and oSL14279/oSL14280 was used to amplify host *fliC* cDNA. Reactions were prepared in 384-well clear/white PCR plates (BioRad), and measurements were performed on a CFX384 RealTime PCR Detection System (BioRad) using the following thermal cycling parameters: polymerase activation and DNA denaturation (98 °C for 2.5 min), 35 cycles of amplification (98 °C for 10 s, 62 °C for 20 s). For each sample, Cq values were normalized to that of *rrsA* (reference housekeeping gene). Then, the normalized Cq values were compared to the normalized Cq value of *fliC* in the control strain (sSL3868, knock-in of *cmR* downstream of *tldR* in BIDMC93), to obtain relative expression levels, such that a value of one is equal to that of the control and higher values indicate higher expression levels.

## Data availability

Next-generation sequencing data are available in the National Center for Biotechnology Information (NCBI) Sequence Read Archive (BioProject Accession: PRJNA1029663) and the Gene Expression Omnibus (GSE245749). The published genome used for ChIP-seq analyses was obtained from NCBI (GenBank: NC_000913.3). The published genomes used for bioinformatics analyses were obtained from NCBI (**Supplementary Table 1**). Datasets generated and analyzed in the current study are available from the corresponding author upon reasonable request.

## Code availability

Custom scripts used for bioinformatics, TAM library analyses, and ChIP-seq data analyses are available upon request.

**Supplementary Information** includes Supplementary Figures 1–6, Supplementary Tables 1–5, and Supplementary Datasets 1–3.

## ACKNOWLEDGMENTS

We thank S.R. Pesari and Z. Akhtar for laboratory support, A. Bernheim for helpful discussions, F. Tesson, A. Bernheim, A.M. Earl, and D. Gray for kindly sharing *E. coli* and *Enterobacter* strains, C. Lu for Covaris sonicator access, L.F. Landweber for qPCR instrument access, and the JP Sulzberger Columbia Genome Center for NGS support. S.T. was supported by a Medical Scientist Training Program grant (5T32GM145440-02) from the NIH. M.W.G.W. was supported by a National Science Foundation Graduate Research Fellowship. C.M. was supported by NIH Postdoctoral Fellowship F32 GM143924-01A1. S.H.S. was supported by NSF Faculty Early Career Development Program (CAREER) Award 2239685, a Pew Biomedical Scholarship, an Irma T. Hirschl Career Scientist Award, and a generous startup package from the Columbia University Irving Medical Center Dean’s Office and the Vagelos Precision Medicine Fund.

## AUTHOR CONTRIBUTIONS

T.R.W., C.M., and S.H.S. conceived and designed the project. T.R.W. performed all bioinformatics experiments and aided in the design of experimental assays. F.T.H. performed plasmid interference, ChIP-seq, and RFP repression assays. M.W.G.W. designed and generated *E. coli* strains and plasmids for RFP repression assays and fragments for *Enterobacter* recombineering. S.T. performed and analyzed RNA-seq and RIP-seq experiments. E.R. cultured *Enterobacter* strains, extracted RNA for RNA-seq, and performed RT-qPCR and recombineering experiments. C.M. performed preliminary TnpB bioinformatics and neighborhood analyses, together with H.C.L., and helped design ChIP-seq and RFP repression assays. T.R.W. and S.H.S. discussed the data and wrote the manuscript, with input from all authors.

## EXTENDED DATA FIGURES

**Extended Data Figure 1.**
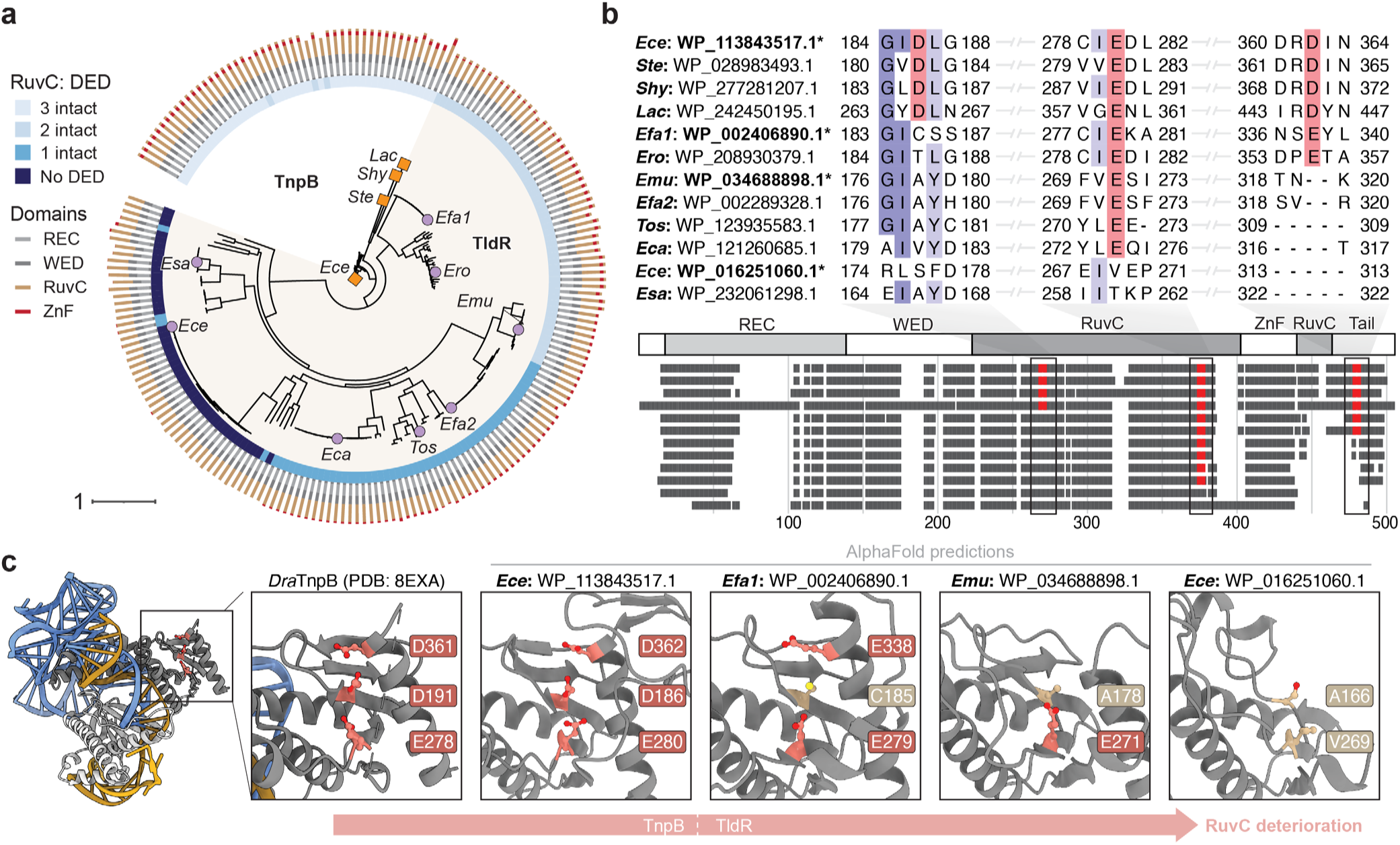
Phylogeny and RuvC nuclease domain analysis of *oppF*-associated TldRs. **a,** Phylogenetic tree of *oppF-*associated TldR proteins from Figure 2a, together with closely related TnpB proteins that contain intact RuvC active sites. The rings indicate RuvC DED active site intactness (inner) and TldR/TnpB domain composition (outer). Homologs marked with an organge square (TnpB) or purple circle (TldR) were tested in heterologous experiments. **b,** Multiple sequence alignment of representative TnpB and TldR sequences from **a**, highlighting deterioration of RuvC active site motifs and loss of the C-terminal Zinc-finger (ZnF)/RuvC domain. **c,** Empirical (*Dra*TnpB) and predicted AlphaFold structures of TnpB and TldR homologs marked with an asterisk in **b**, showing progressive loss of the active site catalytic triad.

**Extended Data Figure 2.**
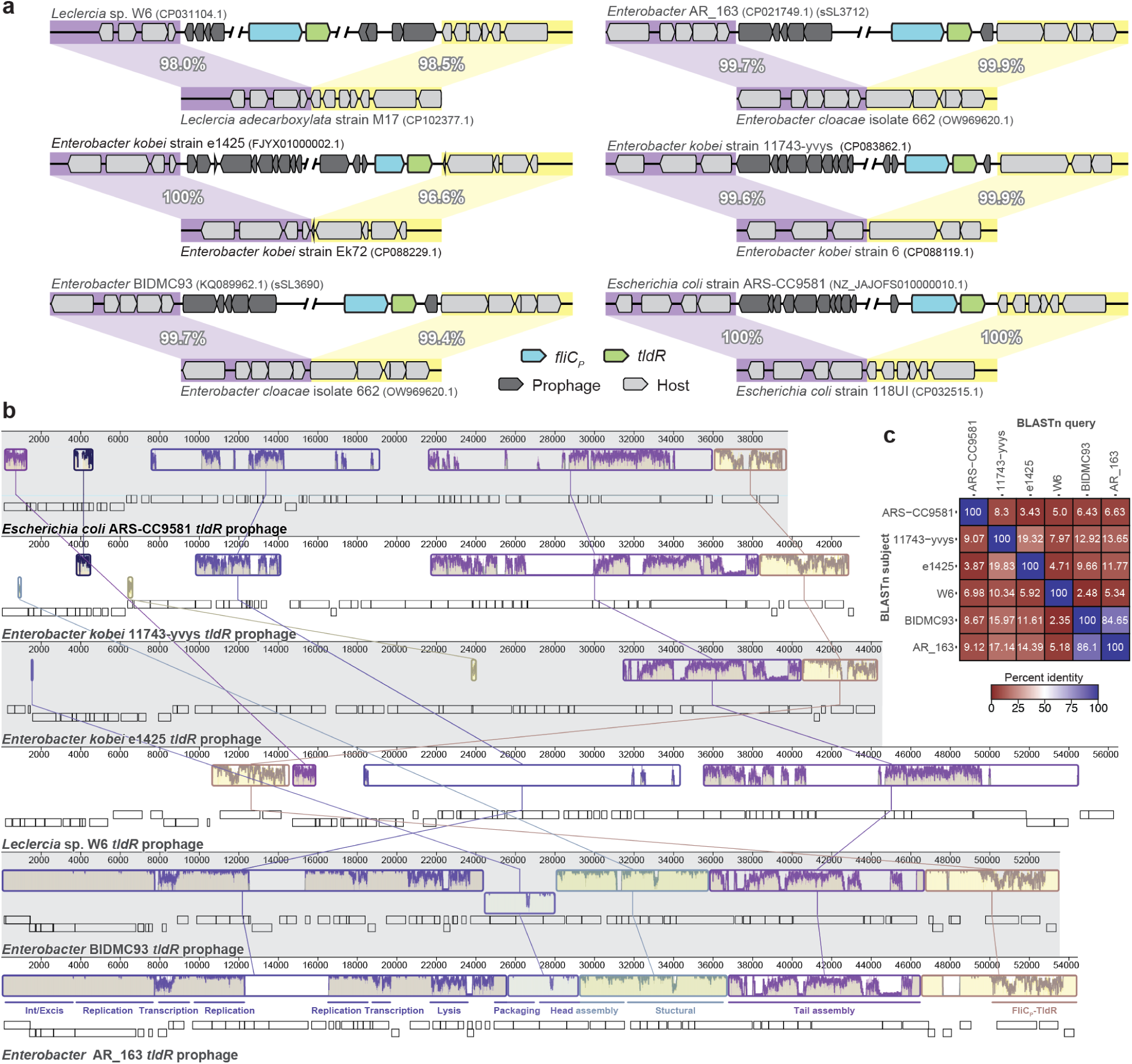
Diverse prophages encode *fliC_P_*-associated *tldR* genes. **a,** Genomic architecture of representative prophage elements whose boundaries could be identified by comparing to closely related isogenic strains. In each example, the prophage-containing strain is shown above the prophage-less strain, with species/strain names and NCBI genomic accession IDs indicated. Sequences flanking the left (5′) and right (3′) ends are highlighted in purple and yellow, respectively, together with their percentage sequence identifies calculated using BLASTn. **b,** Alignment of distinct prophage elements, constructed using Mauve. Empty boxes represent open reading frames, and windows show sequence conservation for regions compared between prophage genomes with lines. Putative gene functions are shown below sequence conservation windows for the *fliC_P_*-*tldR*-encoding prophage from *Enterobacter* AR_163 (bottom). **c,** DNA sequence identities between the prophages in **a**, calculated with BLASTn. Identities were calculated as total matching nucleotides across the two genomes being compared, divided by the length of the query prophage genome.

**Extended Data Figure 3.**
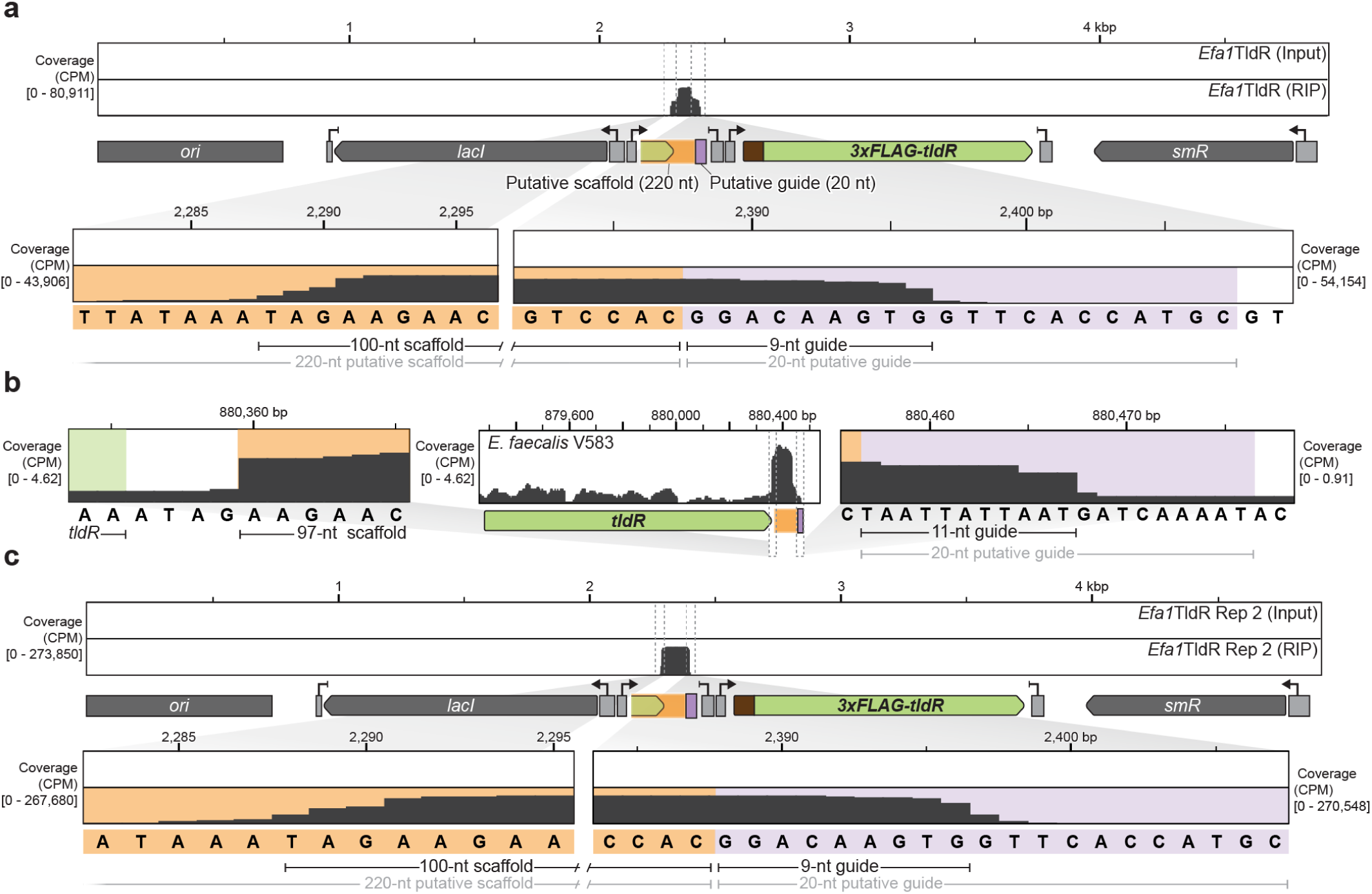
RIP-seq reveals that some *oppF*-associated TldR proteins use short, 9–11-nt guides. **a,** RNA immunoprecipitation sequencing (RIP-seq) data for an *oppF-*associated TldR homolog from *Enterococcus faecalis* (*Efa1*TldR) reveals the boundaries of a mature gRNA containing a 9-nt guide sequence. Reads were mapped to the TldR-gRNA expression plasmid; an input control is shown. **b,** Published RNA-seq data for *Enterococcus faecalis* V583 reveals similar gRNA boundaries, including an ∼11-nt guide. **c,** RIP-seq data as in **a** for a second biological replicate of *Efa1*TldR, further corroborating the observed ∼9–11-nt guide length.

**Extended Data Figure 4.**
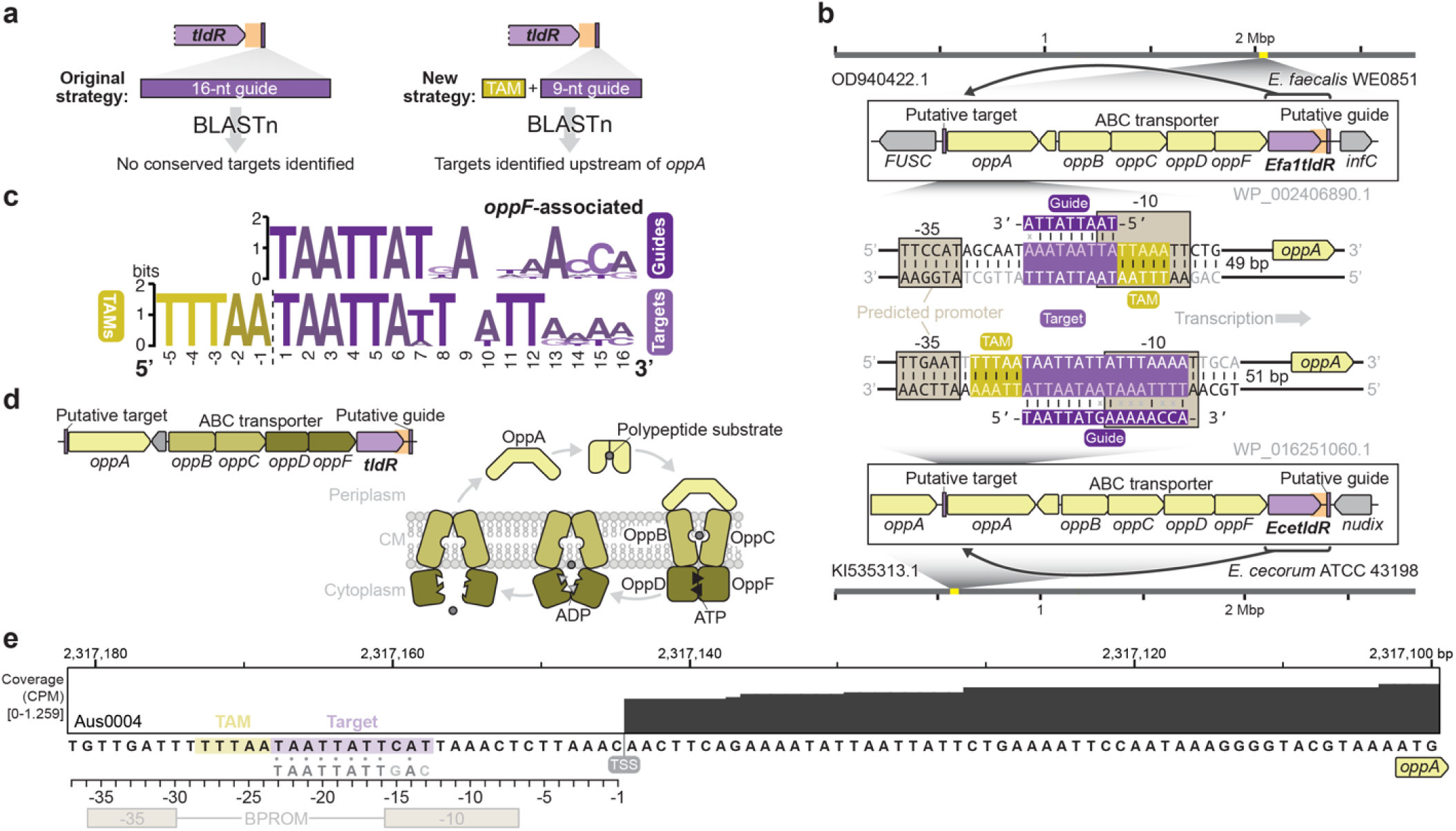
*oppF*-associated TldRs target conserved genomic sequences that overlap with promoter elements driving *oppA* expression. **a,** Schematic of original (left) and new (right) search strategy to identify putative targets of gRNAs used by *oppF*-associated TldRs. Key insights resulted from the use of TAM and a shorter, 9-nt guide. **b,** Analysis of the guide sequence from the *Efa1*TldR-associated gRNA in **Extended Data Figure 3** revealed a putative genomic target near the predicted promoter of *oppA* encoded within the same ABC transporter operon immediately adjacent to the *tldR* gene. The magnified schematics at the bottom show the predicted TAM and gRNA-target DNA base-pairing interactions for two representatives (*Efa1*TldR and *Ece*TldR), in which the gRNAs target opposite strands. Promoter elements predicted with BPROM are shown as brown squares. **c,** WebLogos of predicted guides and genomic targets associated with diverse *oppF*-associated TldRs highlighted in **Supplementary Figure S3a**. **d,** Schematic of the *oppF*-*tldR* genomic locus (left) alongside the predicted function of OppA as a solute binding protein that facilitates transport of polypeptide substrates from the periplasm to the cytoplasm, in complex with the remainder of the ABC transporter apparatus. CM, cell membrane. **e,** Published RNA-seq data for *Enterococcus faecium* AUS0004^46^, highlighting the *oppA* transcription start site (TSS). The predicted gRNA guide sequence (grey) is shown beneath the putative TAM (yellow) and target (purple) sequences, with guide-target complementarity represented by grey circles.

**Extended Data Figure 5.**
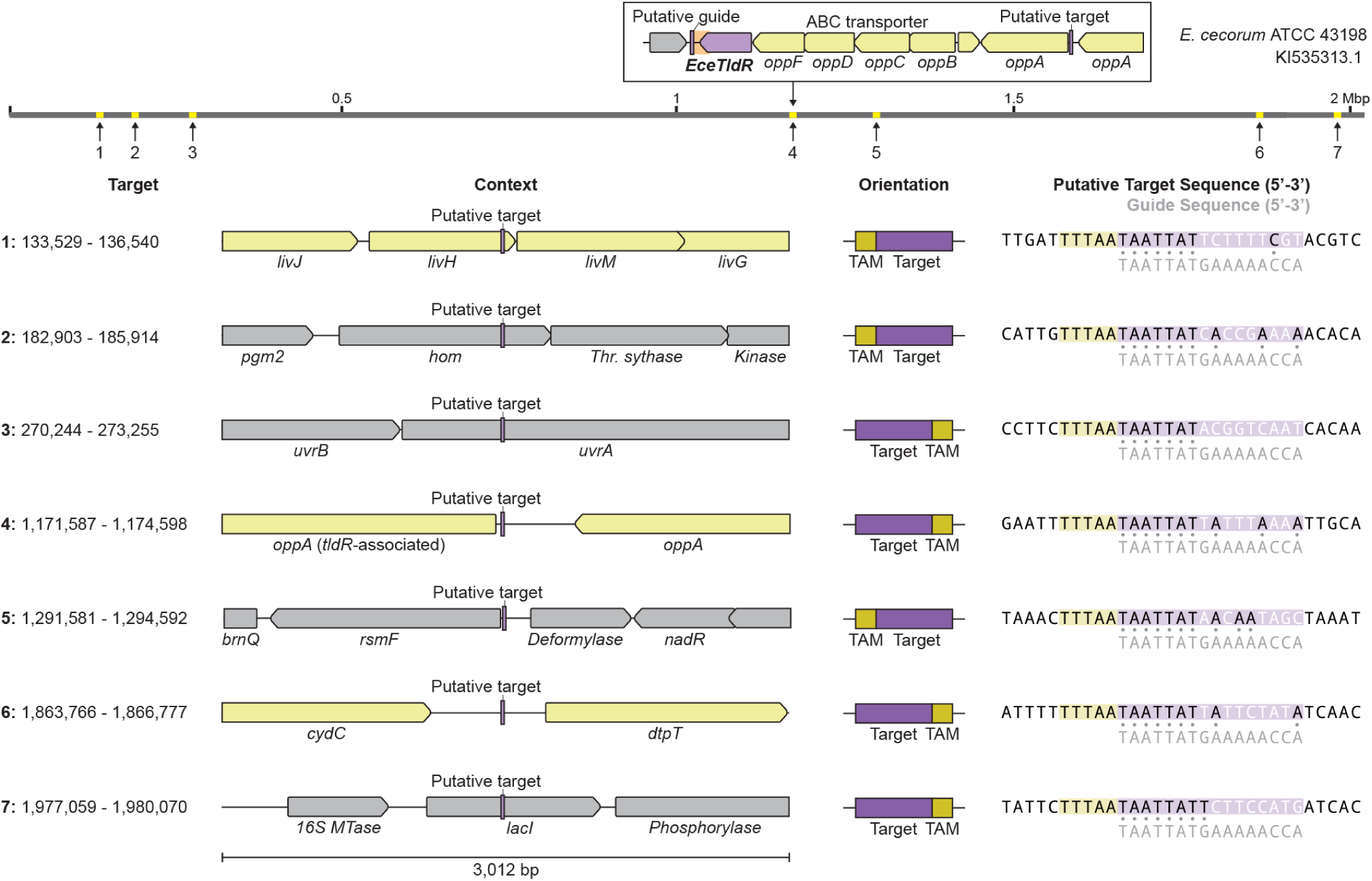
*oppF*-associated TldR homologs may target additional sites across the genome. Schematic of *Enterococcus cecorum* genome and inset showing the *oppF*-*tldR* locus (top), with additional putative targets of the gRNA, other than the *oppA* promoter, numbered and highlighted in yellow along the genomic coordinate. A magnified view for each numbered target is shown below, with TAMs in yellow, prospective targets in purple, and TldR gRNA guide sequences in grey. Grey circles (right) represent positions of expected guide-target complementarity.

**Extended Data Figure 6.**
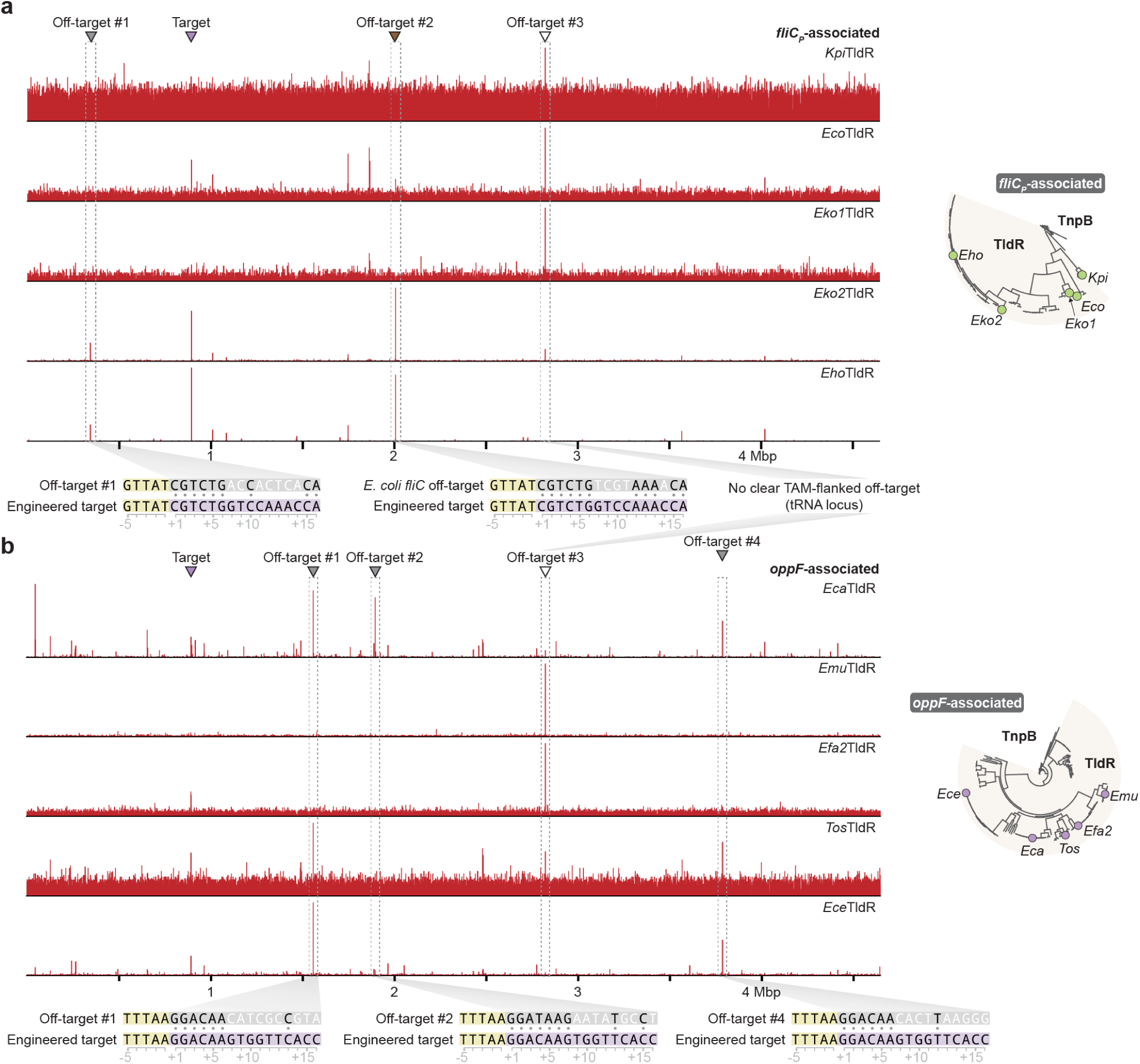
Genome-wide binding data from ChIP-seq experiments suggests a high mismatch tolerance for some TldR homologs. **a,** Genome-wide ChIP–seq profiles for the indicated *fliC_P_*-associated TldR homologs, normalized to the highest peak within each dataset. The magnified insets at the bottom show the off-target sequences (grey) compared to the intended (engineered) on-target sequence (purple), with TAMs in yellow. Off-target #3 has no clear TAM-flanked off-target sequence but is intriguingly located at a tRNA locus, and binding was observed for diverse *fliC_P_*- and *oppF-*associated TldRs that recognized distinct TAMs. The phylogenetic tree at right indicates the relatedness of the tested and labeled homologs. **b,** Results for the indicated *oppF*-associated TldR homologs, shown as in **a**.

**Extended Data Figure 7.**
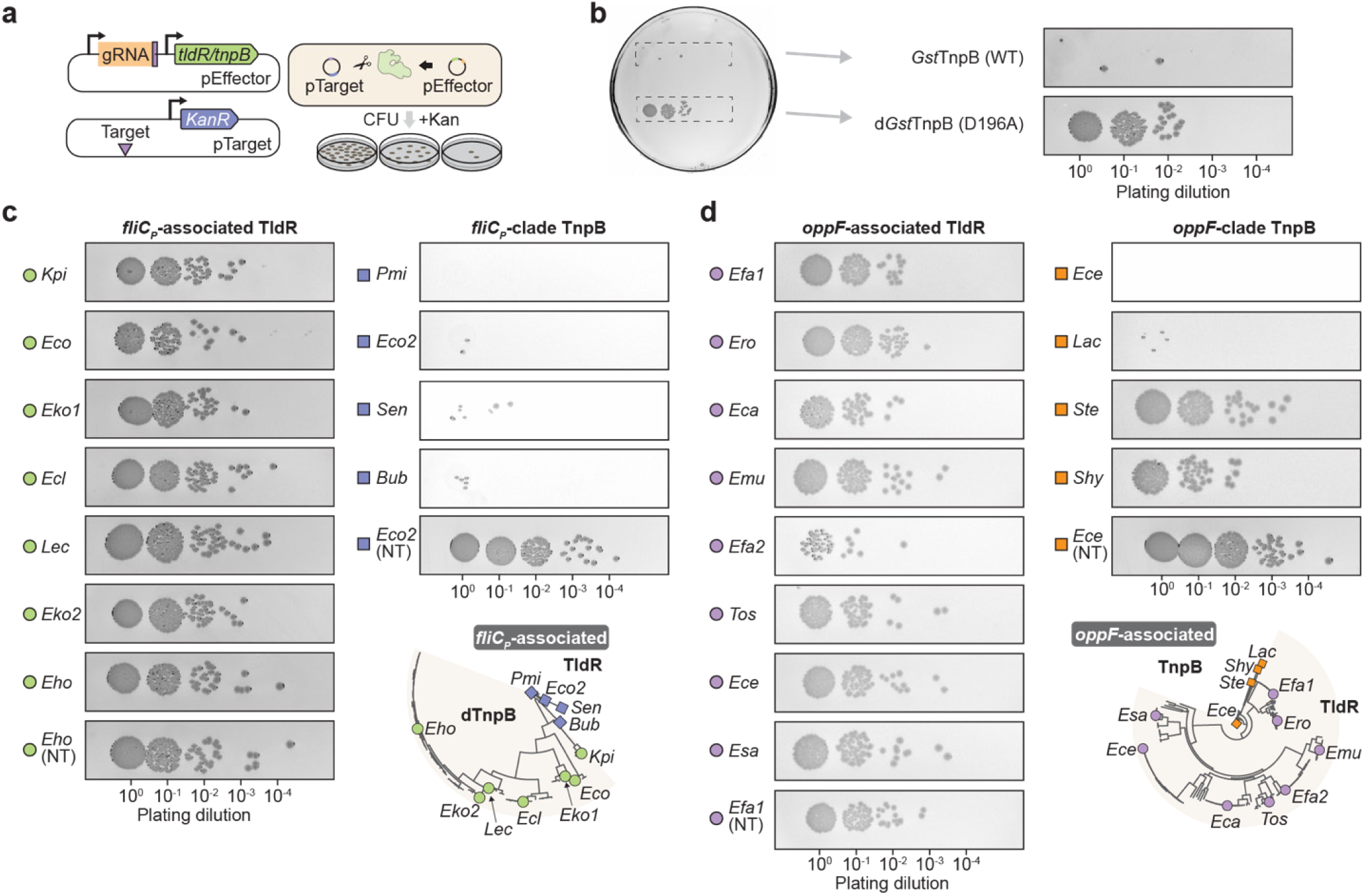
Plasmid interference assays confirm that TldR homologs lack detectable nuclease activity. **a,** Schematic of *E. coli*-based plasmid interference assay using pEffector and pTarget. **b,** Representative dilution spot assays for *Gst*TnpB3 and synthetically inactivated RuvC mutant (D196A), showing the entire plate (left) and the magnified area of plating. Transformants were serially diluted, plated on selective media, and cultured at 37 °C for 16 h. Colony visibility was enhanced by inverted the colors and increasing contrast/brightness. **c,** Dilution spot assays for the indicated *fliC*-associated TldR homologs (left) and closely related TnpB homologs (right). Non-targeting (NT) gRNA controls are shown at the bottom, and the phylogenetic tree indicates the relatedness of the tested proteins. **b,** Results for the indicated *oppF*-associated TldR and TnpB homologs, shown as in **c**.

**Extended Data Figure 8.**
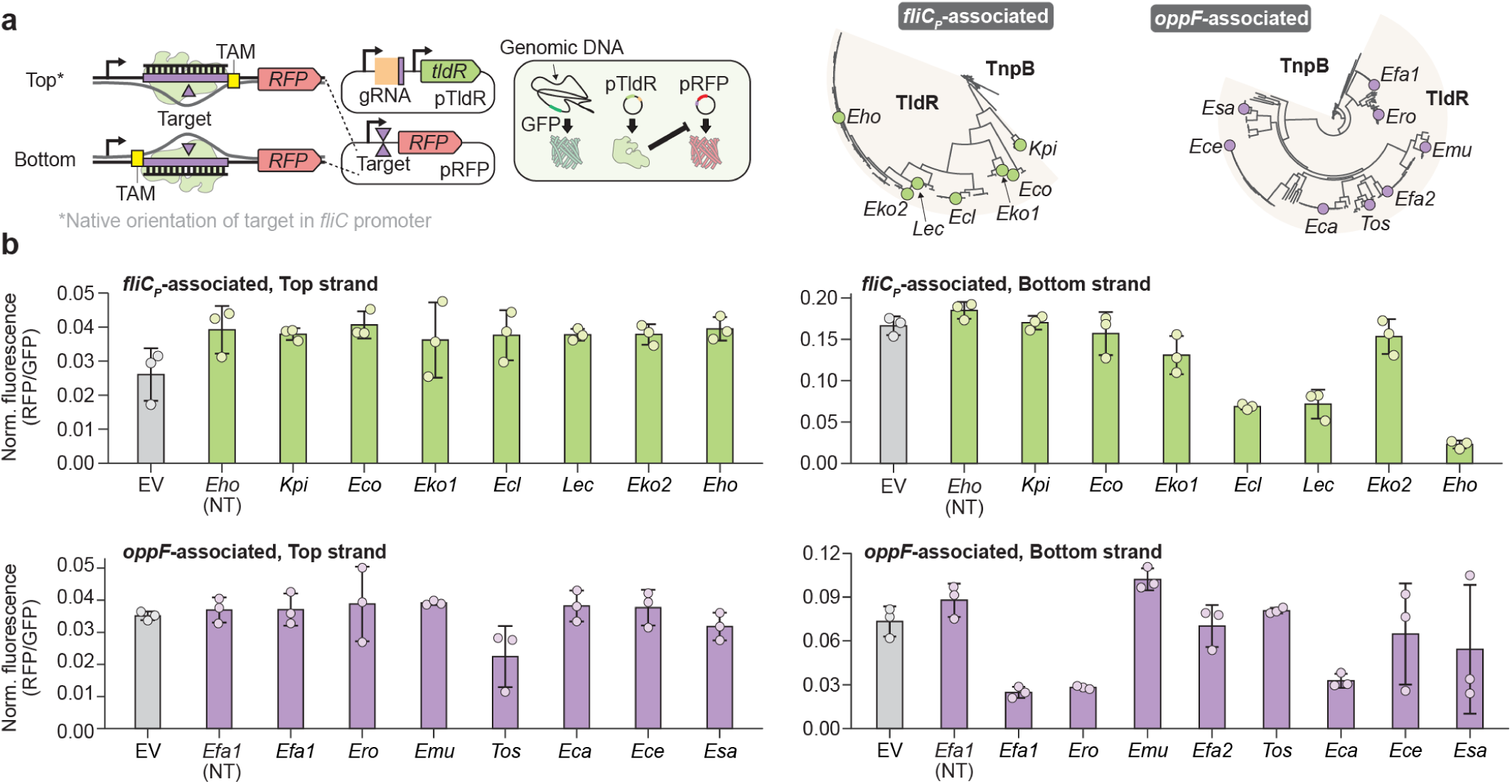
RFP repression assays reveal variable abilities of TldR homologs to block transcription elongation. **a,** Schematic of RFP repression assay adapted from Figure 4g (left), in which gRNAs were designed to target either the top or bottom strand within the 5′ UTR of *RFP*, downstream of the promoter. The phylogenetic trees (right) indicate the relatedness of the tested and labeled homologs. **b,** Bar graph plotting normalized RFP fluorescence for the indicated conditions and TldR homologs. EV, empty vector; NT, non-targeting guide. Bars indicate mean ± s.d. (n = 3).

**Extended Data Figure 9.**
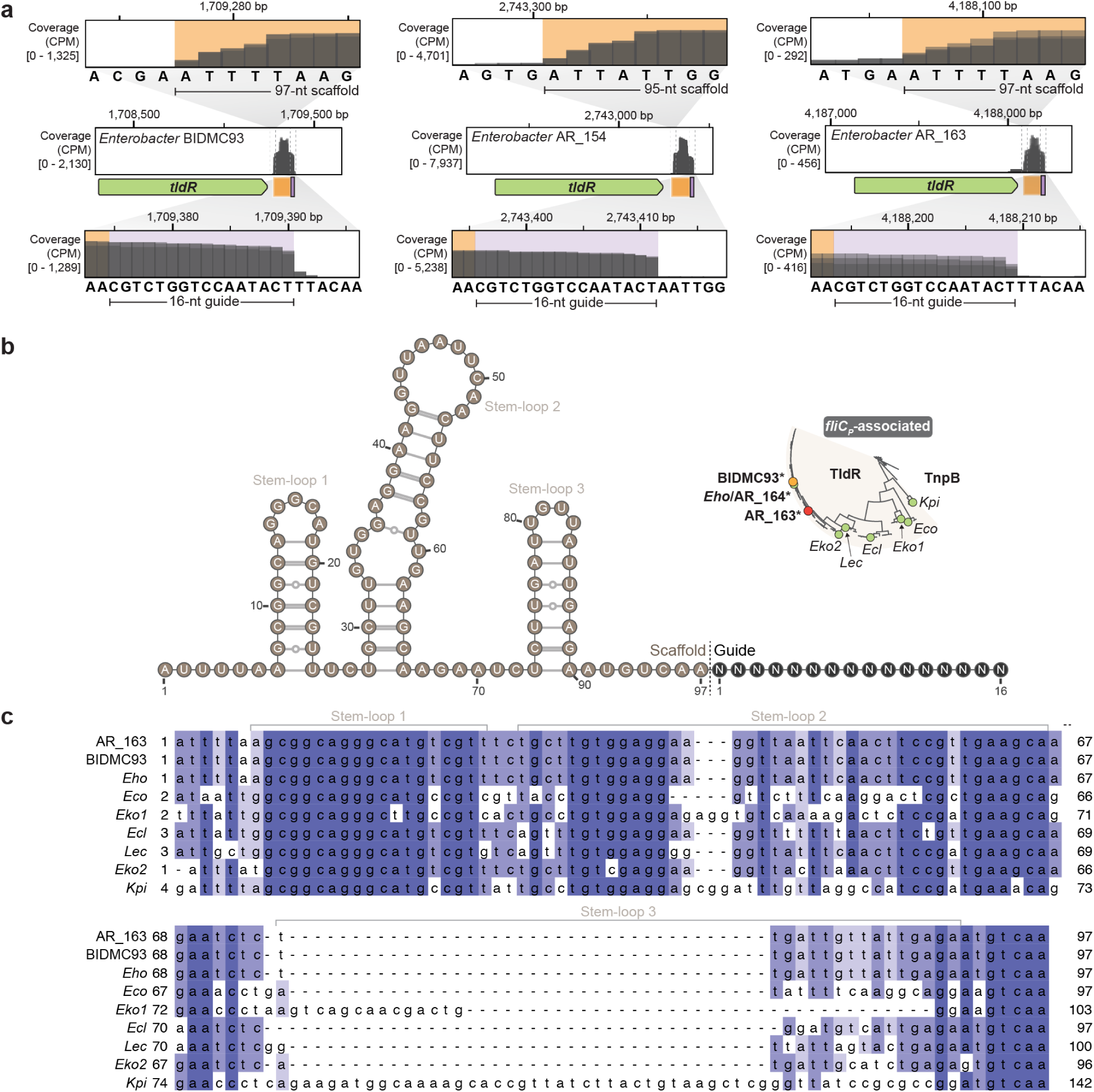
*Enterobacter* RNA-seq data confirm the native expression of gRNAs from *fliC_P_*-*tldR* loci. **a,** RNA-seq read coverage from three *Enterobacter* strains that natively encode *fliC_P_*-*tldR* loci, revealing clear peaks associated with mature gRNAs containing ∼95–97-nt scaffolds and 16-nt guides. Data from three biological replicates are overlaid. **b,** Predicted secondary structure and sequence of the gRNA associated with *Eho*TldR. **c,** Multiple sequence alignment of the DNA encoding gRNA scaffold sequences for representative *fliC_P_*-associated TldRs, with conserved positions colored in darker blue.

**Extended Data Figure 10.**
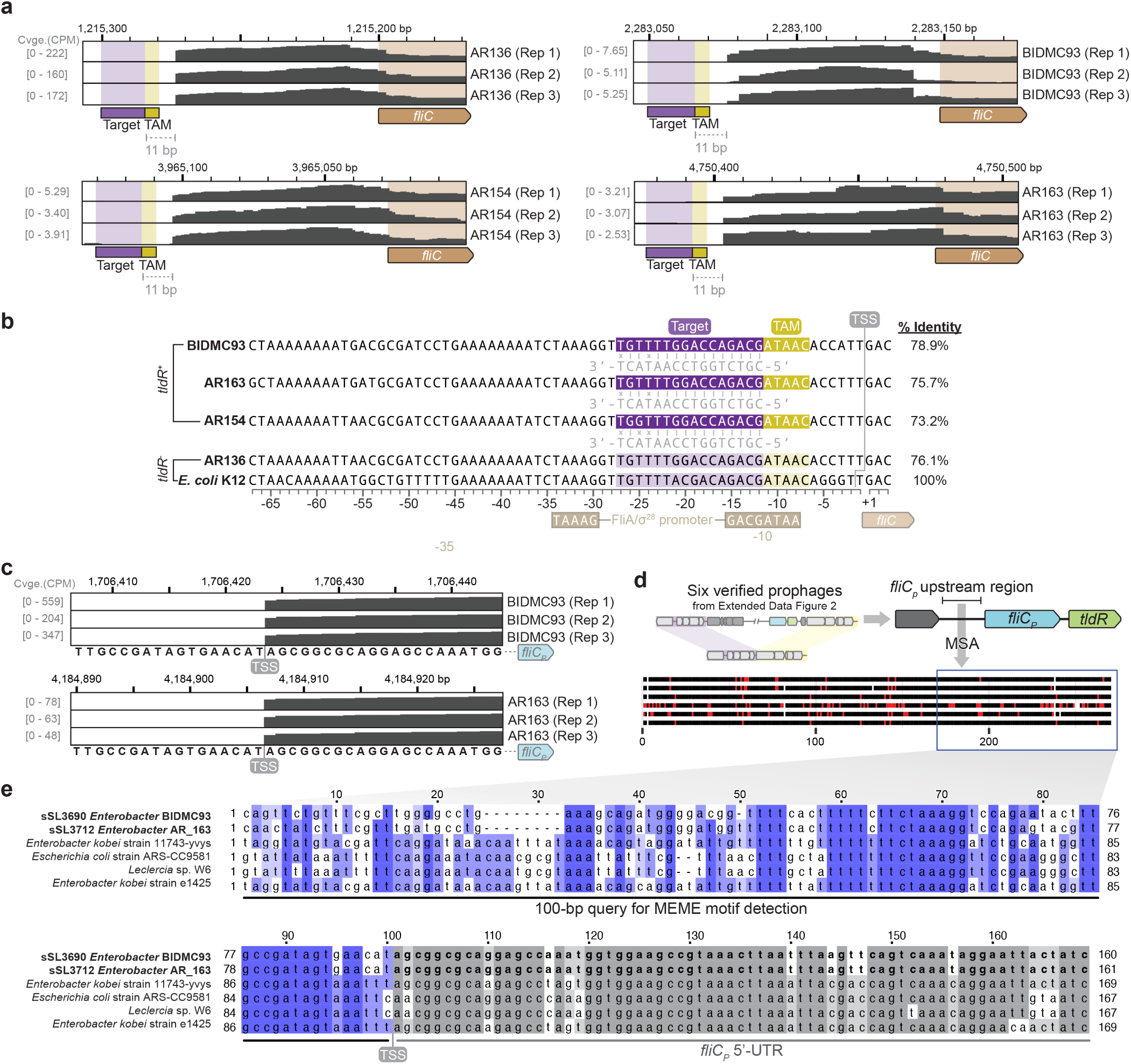
*Enterobacter* RNA-seq data confirm the overlap between TldR-gRNA binding sites and host *fliC* promoters. **a,** RNA-seq read coverage in the host *fliC* promoter/5′-UTR region for four *Enterobacter* strains, with labeled TAM and target sequences highlighted upstream of the TSS. Strain AR136 (top left) does not encode a *fliC_P_*-*tldR* locus; note the distinct expression levels, measured via relative counts per million (CPM). **b,** Alignment of host *fliC* promoter regions for the strains shown in **a** compared to *E. coli* K12, with percent sequence identities indicated on the right. Reported FliA/σ^28^ promoter elements from *E. coli* K12 are shown below the alignment. **c,** RNA-seq read coverage in the prophage-encoded *fliC_P_* promoter/5′-UTR region for two representative *Enterobacter* strains, confirming the predicted TSS. **d,** Schematic of multiple sequence alignment (MSA) of the promoter region driving *fliC_P_* gene expression, across six verified prophages described in **Extended Data Figure 2**. **e,** Magnified MSA for the indicated region in **d**, highlighting the region that was queried for MEME motif detection.

## SUPPLEMENTARY FIGURES

**Supplementary Figure 1.**
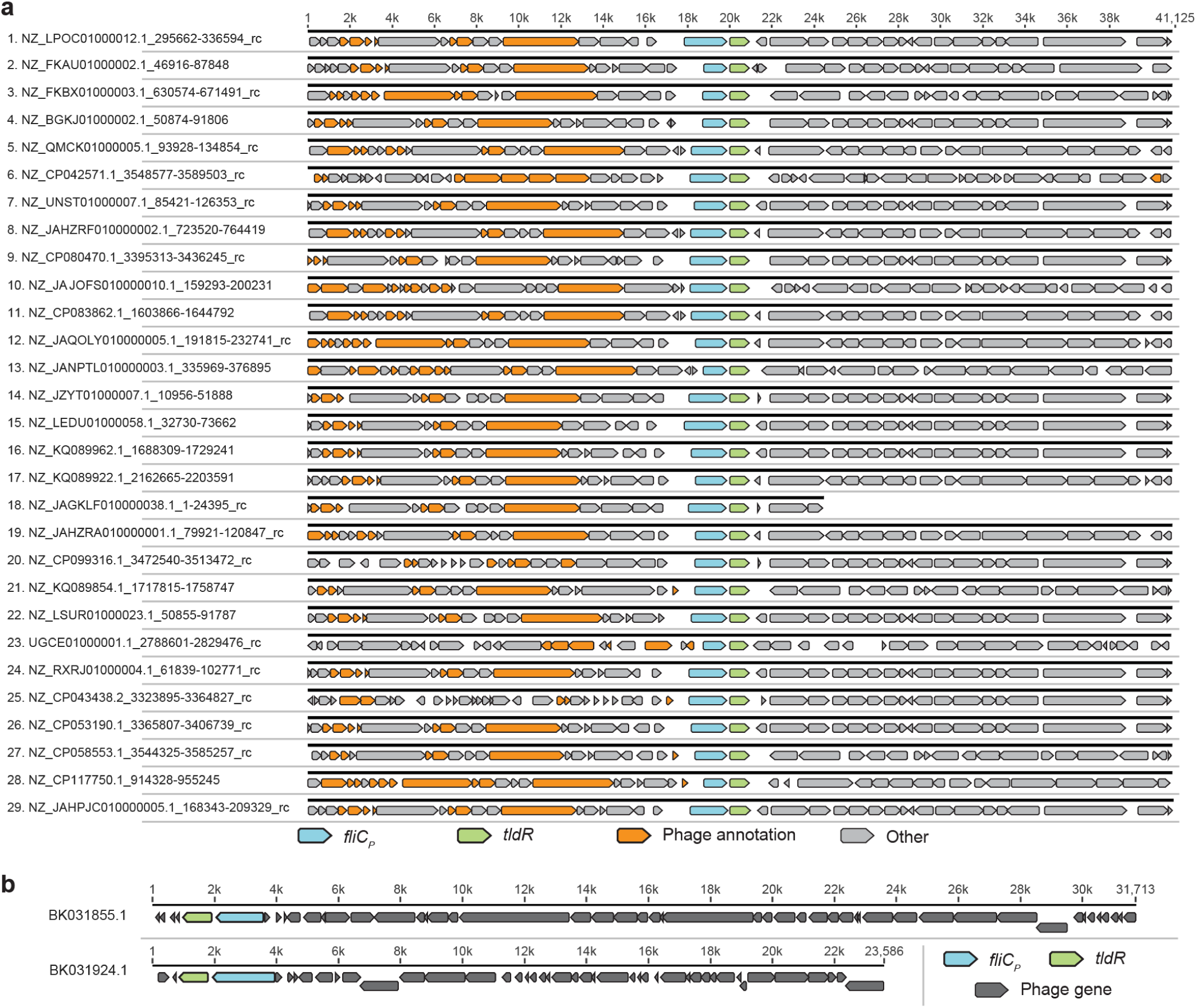
*fliC_P_-tldR* loci are encoded within prophages and phage genomes. **a,** Genetic architecture of a 40 kbp window of bacterial genomes that encode *fliC_P_-tldR* loci (center). *fliC_P_* and *tldR* genes are colored in light blue and green, respectively, and genes with Eggnog annotations containing the word “phage” or “viridae” are colored in orange; all other annotated genes are shown in grey. Each locus is annotated with NCBI accession IDs and genomic coordinates; “_rc” indicates that annotations for the reverse complement sequence are shown. **b**, Two metagenome-assembled phage genomes encode *fliC_P_-tldR* loci. NCBI accessions are shown on the left.

**Supplementary Figure 2.**
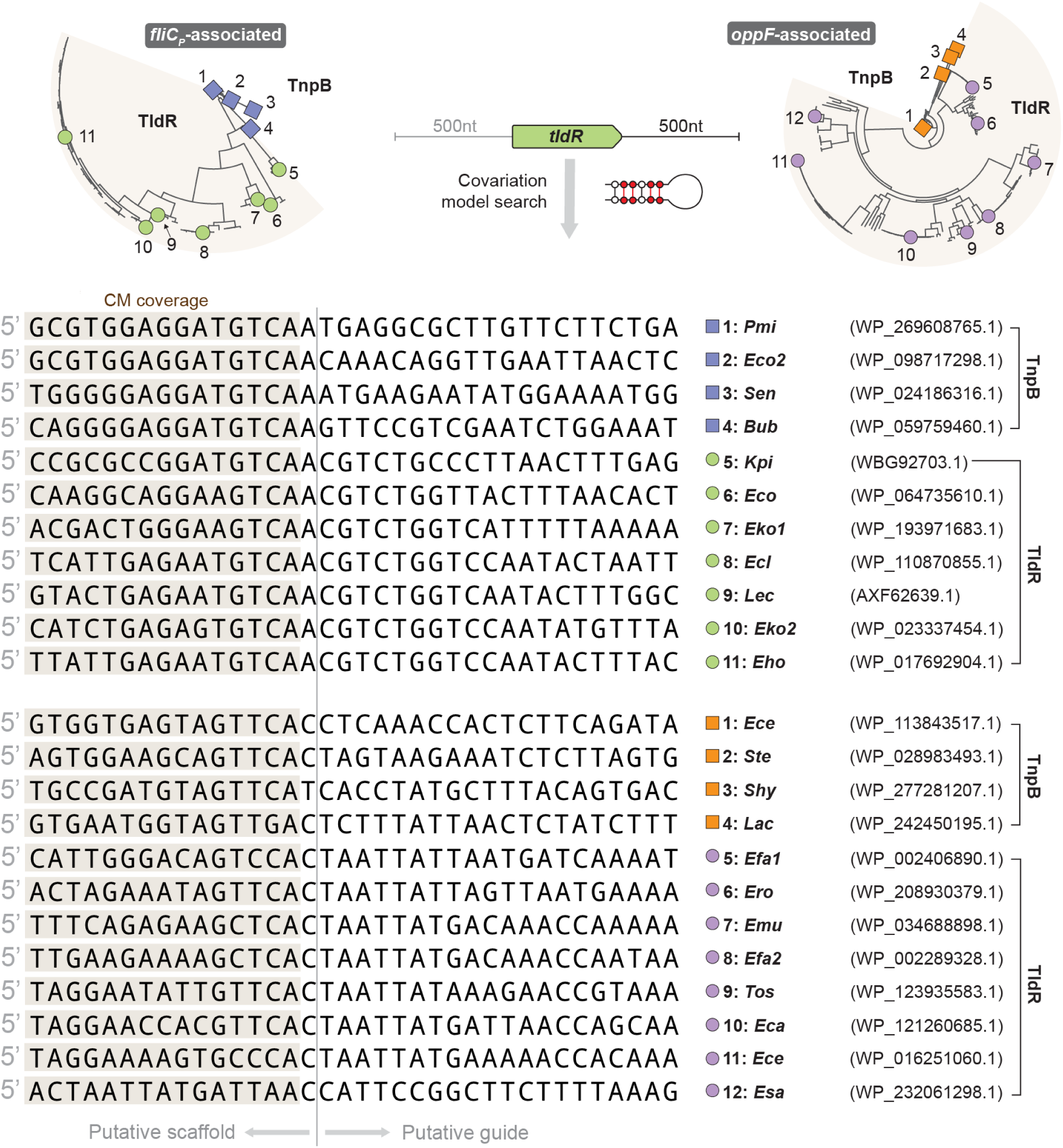
TldR-associated gRNA sequences identified using covariance models. Phylogenetic tree of *fliC-* and *oppF*-associated TldR homologs alongside related TnpB proteins (top), and scaffold/guide junctions for putative TldR-associated gRNAs identified using covariance models (bottom). Matches to the covariance model are shaded, and protein accession IDs are shown at the right.

**Supplementary Figure 3.**
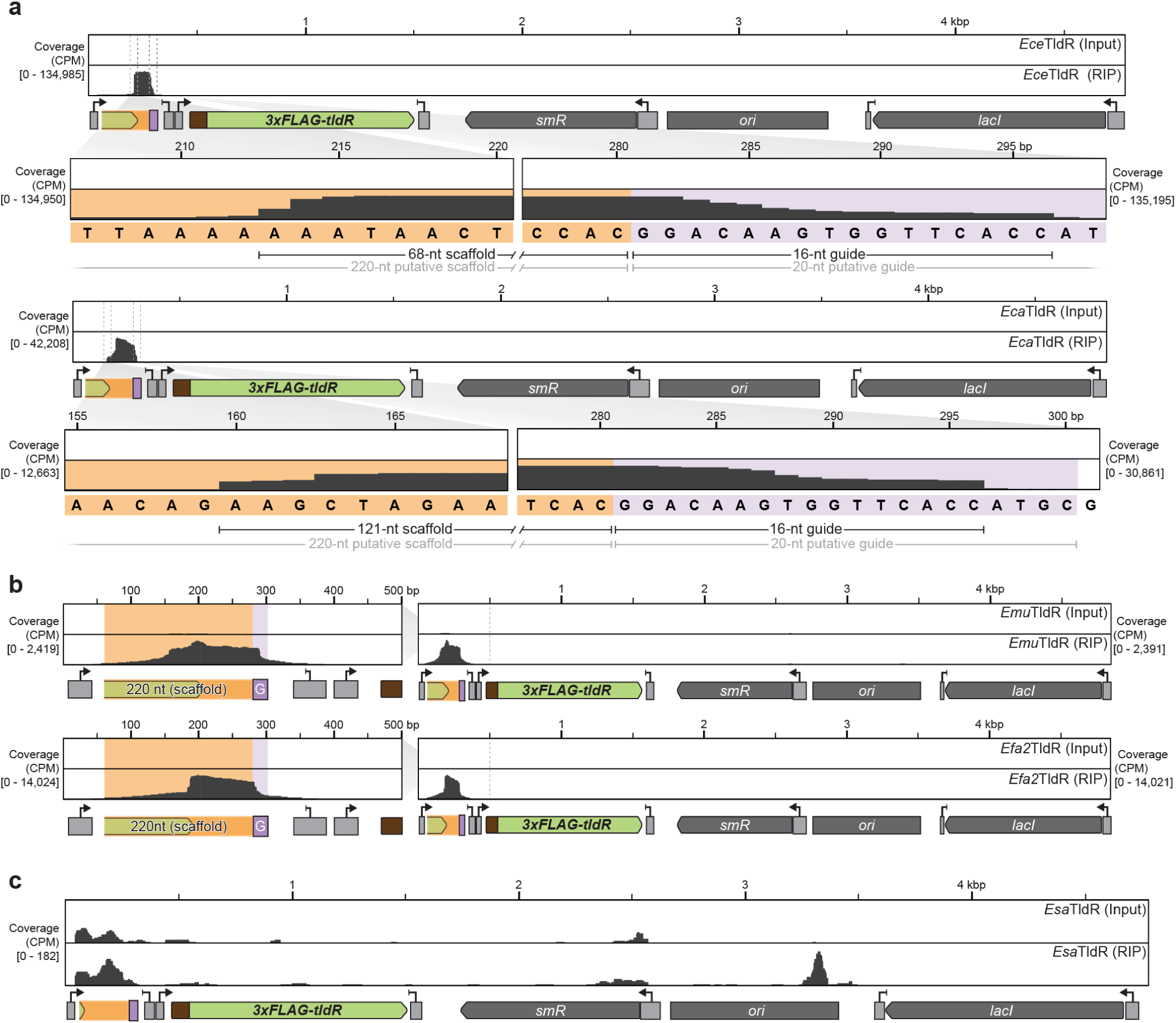
RIP-seq data for additional *oppF*-associated TldR proteins reveals variable gRNA substrates. **a,** RNA immunoprecipitation sequencing (RIP-seq) data for *oppF-*associated TldR homologs from *Enterococcus cecorum* (*Ece*TldR) and *Enterococcus casseliflavus* (*Eca*TldR) indicates variable length guide sequences. Reads were mapped to each respective expression plasmid. **b,** RIP-seq data for *Emu*TldR and *Efa2*TldR, shown as in **a**. **c,** RIP-seq data for *Esa*TldR, shown as in **a**. Enrichment for the gRNA region was not observed, relative to the input control.

**Supplementary Figure 4.**
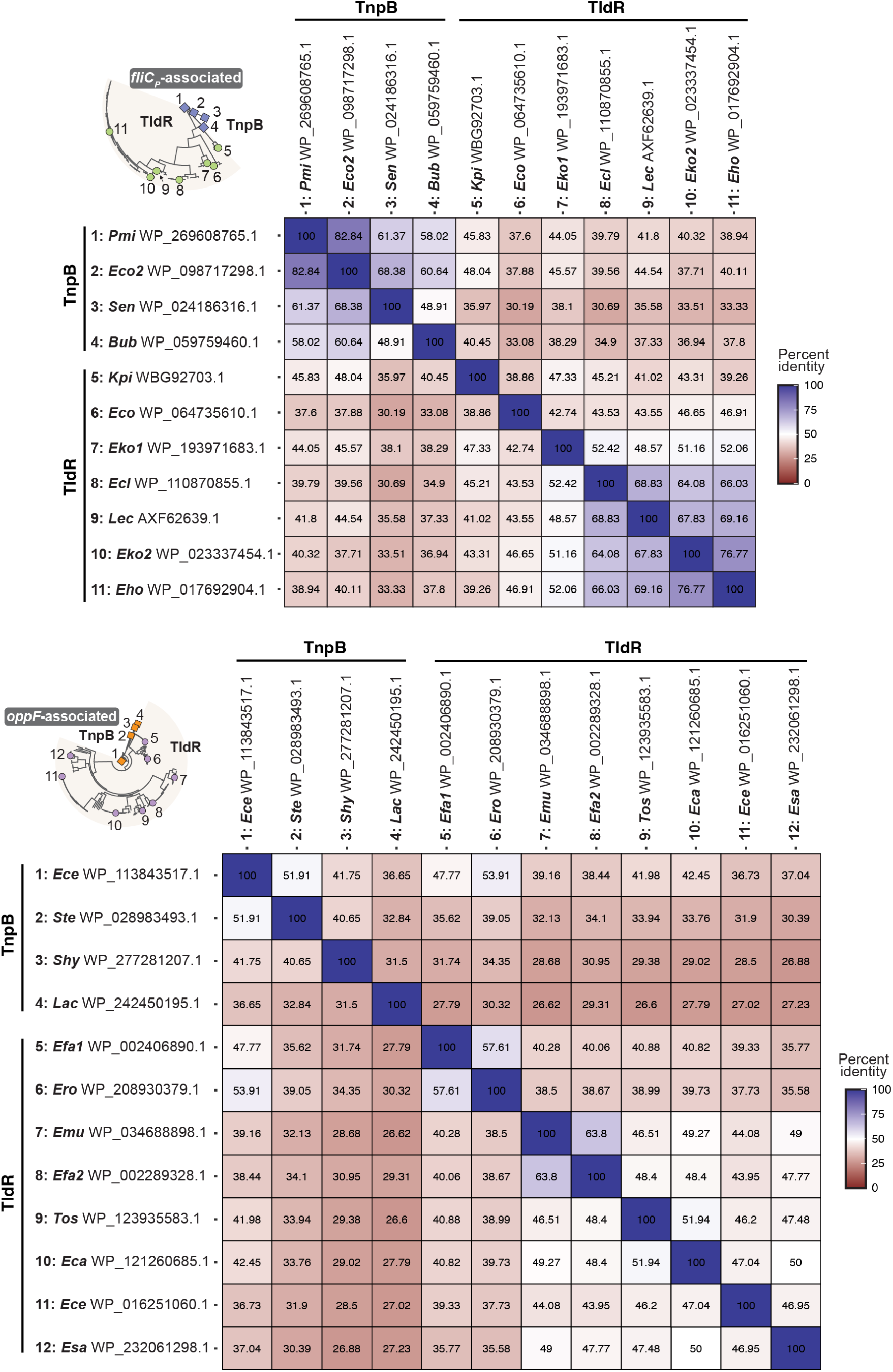
Pairwise identity matrices for representative TldR proteins and related TnpB homologs. Pairwise sequence identities at the amino acid level were calculated for each of the representative TldRs and TnpBs highlighted in **Extended Data Figure 1a**, for *fliC_P_*-associated (top) and *oppF*-associated (bottom) clades.

**Supplementary Figure 5.**
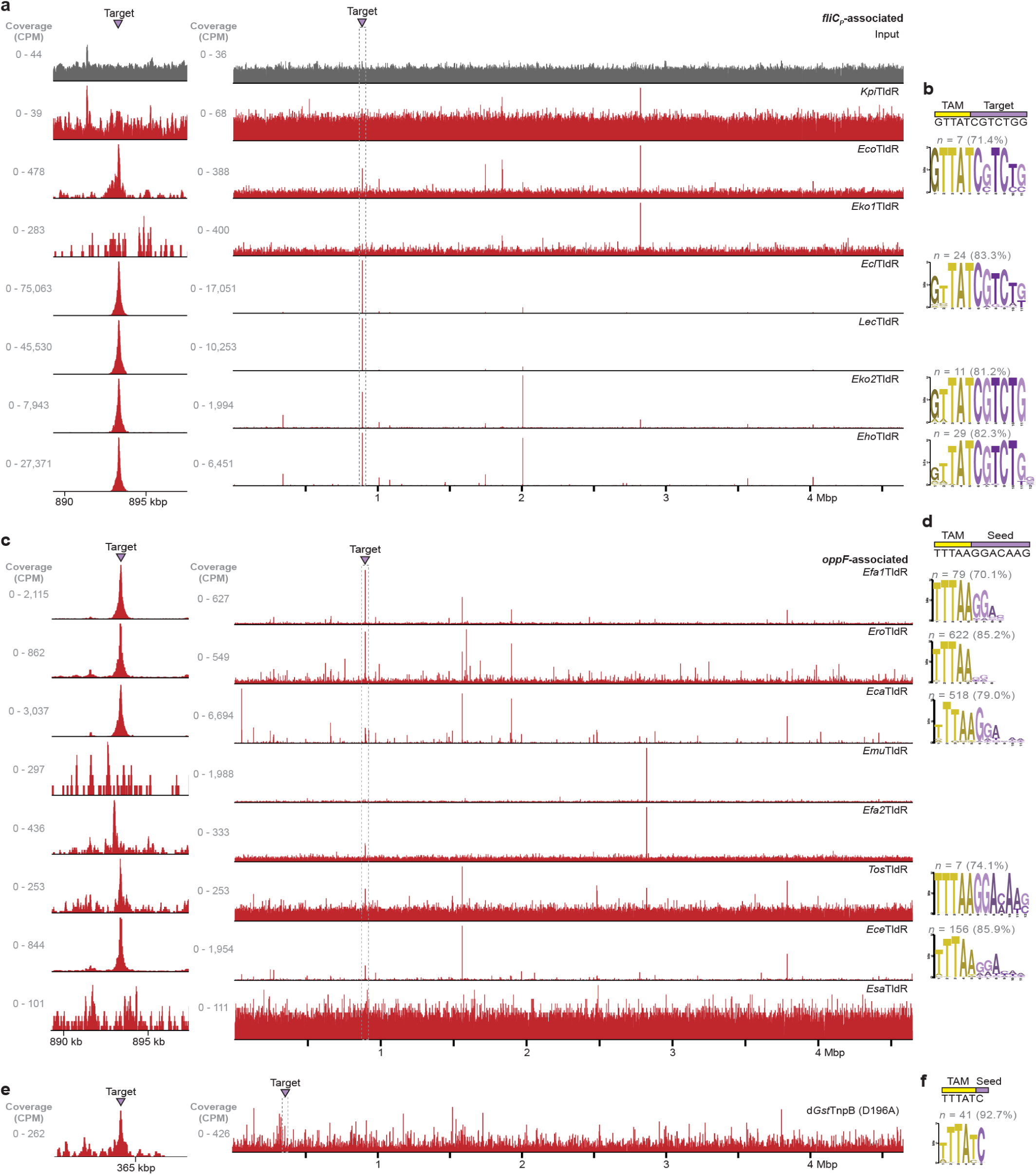
Genome-wide binding data from ChIP-seq experiments for additional TldR homologs. **a,** Genome-wide ChIP–seq profiles for the indicated *fliC_P_*-associated TldR homologs, normalized to the highest peak within each dataset except for the input control (top). The magnified inset at the left shows enrichment at the genomically-integrated, gRNA-matching target site. **b,** Analysis of conserved motifs bound by the indicated TldR homolog in **a** using MEME ChIP, which reveals specificity for the TAM and a ∼6-nt seed sequence. The number of peaks and percentage of total called peaks contributing to each motif is indicated; low occupancy positions were manually trimmed from motif 5′ ends. Motifs are omitted for datasets for which a high-confidence consensus could not be identified. **c,** Genome-wide ChIP–seq profiles for the indicated *oppF*-associated TldR homologs, shown as in **a**. **d,** Analysis of conserved motifs bound by the indicated TldR homolog in **c** using MEME ChIP, shown as in **b**. **e,** Genome-wide ChIP–seq profile for *Gst*TnpB_D196A_, shown as in **a**. **f,** Analysis of conserved motifs bound by *Gst*TnpB_D196A_ in **e** using MEME ChIP, shown as in **b**.

**Supplementary Figure 6.**
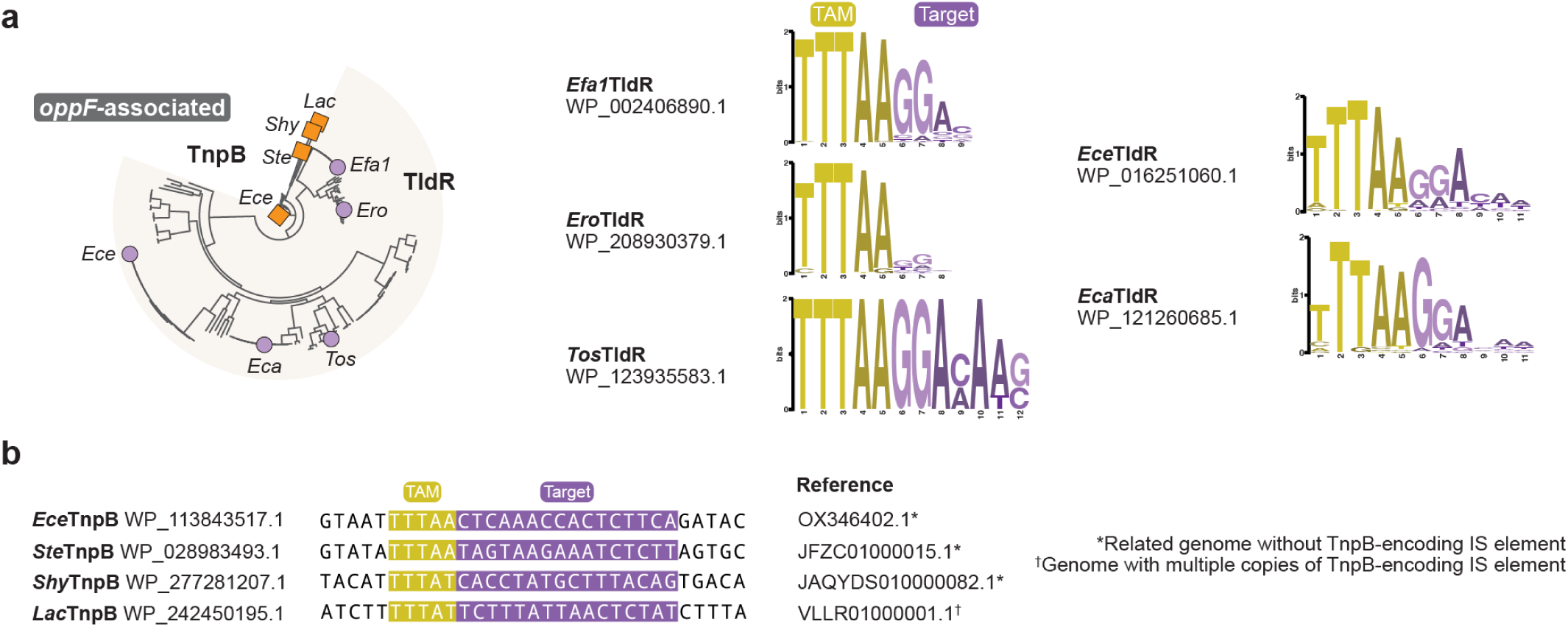
Comparison of TAM specificities for *oppF*-associated TldRs and related TnpBs, determined via ChIP-seq and comparative genomics. **a,** Phylogenetic tree showing the relatedness of labeled *oppF*-associated TldRs and similar TnpB homologs (left), and consensus motifs from TldR homologs using MEME ChIP, replotted from **Supplementary Figure 5**. TAMs and target regions are colored in yellow and purpled, respectively. **b,** Bioinformatically predicted TAMs and target sequences for related TnpB homologs labeled in the tree from **a**. Reference genomes used for comparative genomics analyses to predict the TAM (yellow) and target (purple) are indicated, and harbored either isogenic loci lacking the transposon IS element, or multiple copies of the same IS element.

